# Genetic variation at mouse and human ribosomal DNA influences associated epigenetic states

**DOI:** 10.1101/2021.06.10.447887

**Authors:** Francisco Rodriguez-Algarra, Robert A E Seaborne, Amy F Danson, Selin Yildizoglu, Harunori Yoshikawa, Pui Pik Law, Zakaryya Ahmad, Victoria A Maudsley, Ama Brew, Nadine Holmes, Mateus Ochôa, Alan Hodgkinson, Sarah J Marzi, Madapura M Pradeepa, Matthew Loose, Michelle L Holland, Vardhman K Rakyan

## Abstract

**Background:** Ribosomal DNA (rDNA) displays substantial inter-individual genetic variation in human and mouse. A systematic analysis of how this variation impacts epigenetic states and expression of the rDNA has thus far not been performed.

**Results:** Using a combination of long- and short-read sequencing, we establish that 45S rDNA units in the C57BL/6J mouse strain exist as distinct genetic haplotypes that influence the epigenetic state and transcriptional output of any given unit. DNA methylation dynamics at these haplotypes are dichotomous and life-stage specific: at one haplotype, the DNA methylation state is sensitive to the *in utero* environment, but refractory to post-weaning influences, whereas other haplotypes entropically gain DNA methylation during ageing only. On the other hand, individual rDNA units in human show limited evidence of genetic haplotypes, and hence little discernible correlation between genetic and epigenetic states. However, in both species, adjacent units show similar epigenetic profiles, and the overall epigenetic state at rDNA is strongly positively correlated with total rDNA copy number. Analysis of different mouse inbred strains reveals that in some strains, such as 129S1/SvImJ, rDNA copy number is only approximately 150 copies per diploid genome and DNA methylation levels are <5%.

**Conclusions:** Our work demonstrates that rDNA-associated genetic variation has a considerable influence on rDNA epigenetic state and consequently rRNA expression outcomes. In the future, it will be important to consider the impact of inter-individual rDNA (epi)genetic variation on mammalian phenotypes and diseases.

## Background

The ribosome is one of the fundamental macromolecular complexes in all living cells, enabling translation to occur in the cytoplasm. The mature mammalian 80S ribosome consists of small (40S) and large (60S) subunits, both comprised of a different complement of proteins and RNA. Despite the essential and highly conserved role played by the ribosome, it is now clear that it can be compositionally diverse even within a given individual organism^1^. It has been shown that the protein composition of the ribosome can vary developmentally^1^, and that the ribosomal RNA (rRNA) components display inter- and intra-individual genetic variation^2,3^. Such variation is thought to ultimately influence preferential translation of some mRNAs, i.e., the “ribosome filter hypothesis”^4^.

With respect to rRNA variation in mammals, several studies have reported genetic variation within the 45S rDNA in human and mouse (**Fig. 1A**). The 45S rDNA codes for the 18S rRNA that is incorporated into the 40S subunit, and the 5.8S and 28S rRNAs that are incorporated into the 60S subunit (which additionally contains the 5S rRNA that is coded by the 5S rDNA present at an unlinked genomic location). Because of the multi-copy and multi-allelic nature of 45S rDNA, functional genomic analyses of this source of mammalian genetic variation have been difficult. Indeed, rDNA clusters are typically excluded from genome assemblies, and none of the large-scale genomic analyses of recent years have yielded any insights into rDNA genetic variation. Nonetheless, several smaller studies have noted inter-individual variation at the single nucleotide level and copy number in mouse and human rDNA, and also epigenetic variation in the context of whole organism stress responses^2,5–8^. Furthermore, our previous work showed that mammalian rDNA sequence variation can influence epigenetic states, thereby impacting transcriptional outputs in different biological contexts. However, a systematic analysis of how human and/or mouse rDNA-associated genetic variation influences associated epigenetic states, and rDNA transcriptional output, has thus far not been performed. We therefore had three main aims in this study, specifically to combine long- and short-read sequencing to establish: (i) how single nucleotide genetic variation at the individual rDNA unit level influences epigenetic states and transcriptional outputs; (ii) whether rDNA copy number influences epigenetic states; (iii) if such genetic variation is relevant in examples of mammalian phenotypes.

**Figure 1.**
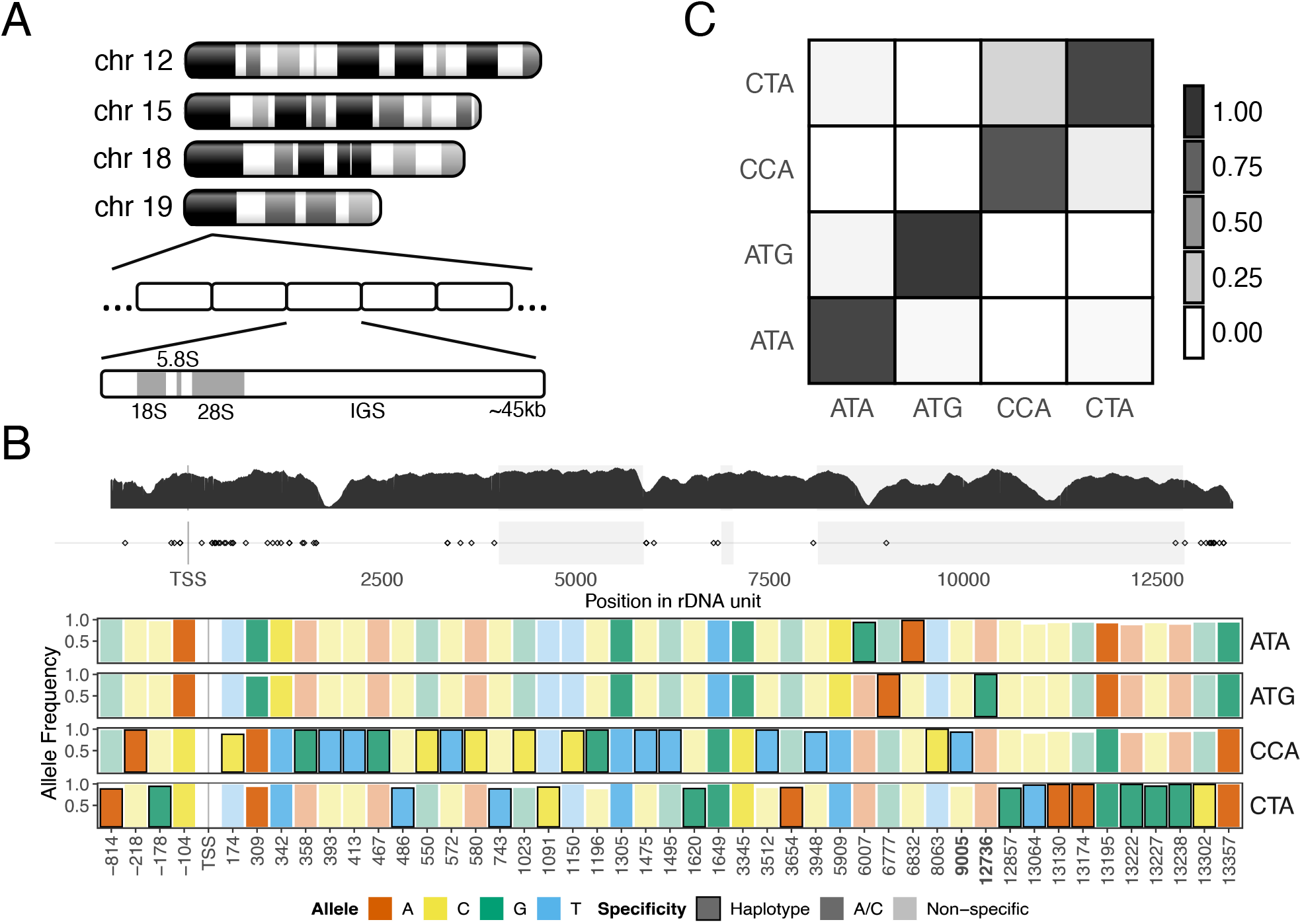
Long range haplotype characterization of rDNA in the C57BL/6J strain. **(A)** Schematic of rDNA in the C57BL/6J mouse strain. Adapted from *Refs 2* and *35* **(B)** rDNA coding unit haplotypes in a C57BL/6J MEF line defined using ultra-long read Nanopore sequencing. The top track shows representative read depth from C57BL/6J kidney short-read whole genome sequencing data, and locations of SNVs cross-validated with Nanopore data. The bottom track shows only SNVs that distinguish rDNA haplotypes. Bold contour denotes variants unique to that specific haplotype. Bars with non-muted colours and no contour indicate positions associated with the A/C haplogroups defined by the variant at position −104. Bolded positions in the x-axis (9005, 12376) are variants within the 28S rRNA and present in mature ribosomes. Haplotypes are denoted by three letters, for the variant nucleotides at positions −104, 8063, and 12736, respectively. 12736 distinguishes the 2 haplotypes with “A” at −104, 8063 distinguishes the 2 haplotypes with “C” at −104 (see **Additional file 3: fig. S2**). Although these three positions combined robustly identify the haplotypes, each individual position is not strictly haplotype-specific. **(C)** Co-localization analysis of rDNA haplotypes in MEF. For Nanopore reads spanning multiple rDNA units, each cell shows the average proportion of units assigned to the corresponding column haplotype in a read, given that the read includes at least one unit of the haplotype indicated in its row.

## Results

### Mouse rDNA exists as distinct genetic haplotypes

Using the C57BL/6J mouse strain, we focussed on a ~15kb region of the rDNA termed the ‘coding unit’ (**Fig. 1A**; the intergenic sequence, IGS, contains high repeat sequence density and was not analysed in detail). Short-read whole genome sequencing (WGS) analysis of four different C57BL/6J mice identified 88 different coding unit intra- and inter-individual single nucleotide variants (SNVs) (**Additional file 1**; **Additional file 2: table S1**; inclusion of indels did not affect SNV allele frequencies and were not explicitly considered further). To achieve a deeper understanding of rDNA genomic architecture, we sequenced a C57BL/6J mouse embryonic fibroblast (MEF) line using ultra-long read whole genome Nanopore technology^9^, obtaining 932,683 reads (N50 ~72 kb), of which 1,760 contained one or more rDNA coding units within a single read (**Additional file 1**; **Additional file 2: table S2**). We confirmed 87/88 short-read rDNA coding unit SNVs in these reads (**Additional file 2: table S1**). Previously, we reported a SNV at position −104 (relative to TSS; **Fig. 1A**) in the C57BL/6J rDNA that is associated with differential promoter methylation: ‘A’ variants at −104 are associated with 30-80% methylation, whereas ‘C’ variants display <25% methylation at the promoter^6,7^. Using −104 as a starting point to explore the possibility of larger haplotypes within the rDNA, analysis of the SNVs throughout the coding unit revealed 4 different rDNA haplotypes, that we term ‘ATA’, ‘ATG’, ‘CCA’, ‘CTA’, in approximately equal proportions (**Fig. 1B**). ATA and ATG are more genetically similar to each other compared to either CCA or CTA (**Fig. 1B**; **Additional file 1**). Independent support for these haplotypes was obtained by pairwise correlation analysis of the relevant haplotype-associated SNVs in the four different short-read kidney WGS datasets (**Additional file 3: fig. S1**; **Additional file 2: table S3**). Analysis of reads containing two or more complete rDNA units revealed that adjacent units tend to correspond to the same rDNA haplotype (**Fig. 1C**). Although further refinement of C57BL/6J rDNA haplotypes may be possible, the analyses below demonstrate that the rDNA haplotypes underlie bona fide molecular differences.

### The impact of rDNA haplotypes on functional genomic outcomes

Nanopore sequencing permits direct assessment of DNA methylation in unamplified DNA. Strikingly, we found that the ATA haplotype displays significant DNA methylation (≳60%) across the length of the coding unit, CCA shows low methylation levels (≲20%), and the other two haplotypes are largely unmethylated (**Fig. 2A**). Analysis of individual reads revealed that individual coding units are either almost completely methylated or unmethylated (**Additional file 3: fig. S2**), and haplotype-specific methylation differences do not extend into the adjoining IGS regions, which are generally hypermethylated (**Additional file 3: fig. S2**). To provide further support for rDNA haplotype-specific methylation using an orthologous method, we analysed 4 different kidney whole-genome bisulfite sequencing (WGBS) and 7 different sperm reduced representation bisulfite sequencing (RRBS) C57BL/6J datasets (11 different mice) (**Additional file 2: table S3**). Short-read methods cannot directly define long-range (epi)genetic patterns, and in our case the major ATA-defining SNV is an ‘A’ at position 6832. However, if the proposed (epi)genomic architecture of the haplotypes is correct, then if we combine the ATA and ATG haplotypes into a single ‘A’ haplogroup, that can be distinguished from the CCA and CTA haplotypes at multiple positions, then in short-read data the allele frequency and DNA methylation state at these positions should be predictable based on the combined individual allele frequencies and DNA methylation levels of ATA and ATG at position 6832. This is indeed what we observe (**Fig. 2B; Additional file 3: fig. S3**). The WGBS and RRBS datasets also demonstrate that whilst relative allele frequencies and DNA methylation show inter-individual variation - in particular, the CCA haplotype is occasionally associated with methylation levels up to ~20% –, it is only the ATA rDNA haplotype that consistently shows substantial methylation *in vivo*.

**Figure 2.**
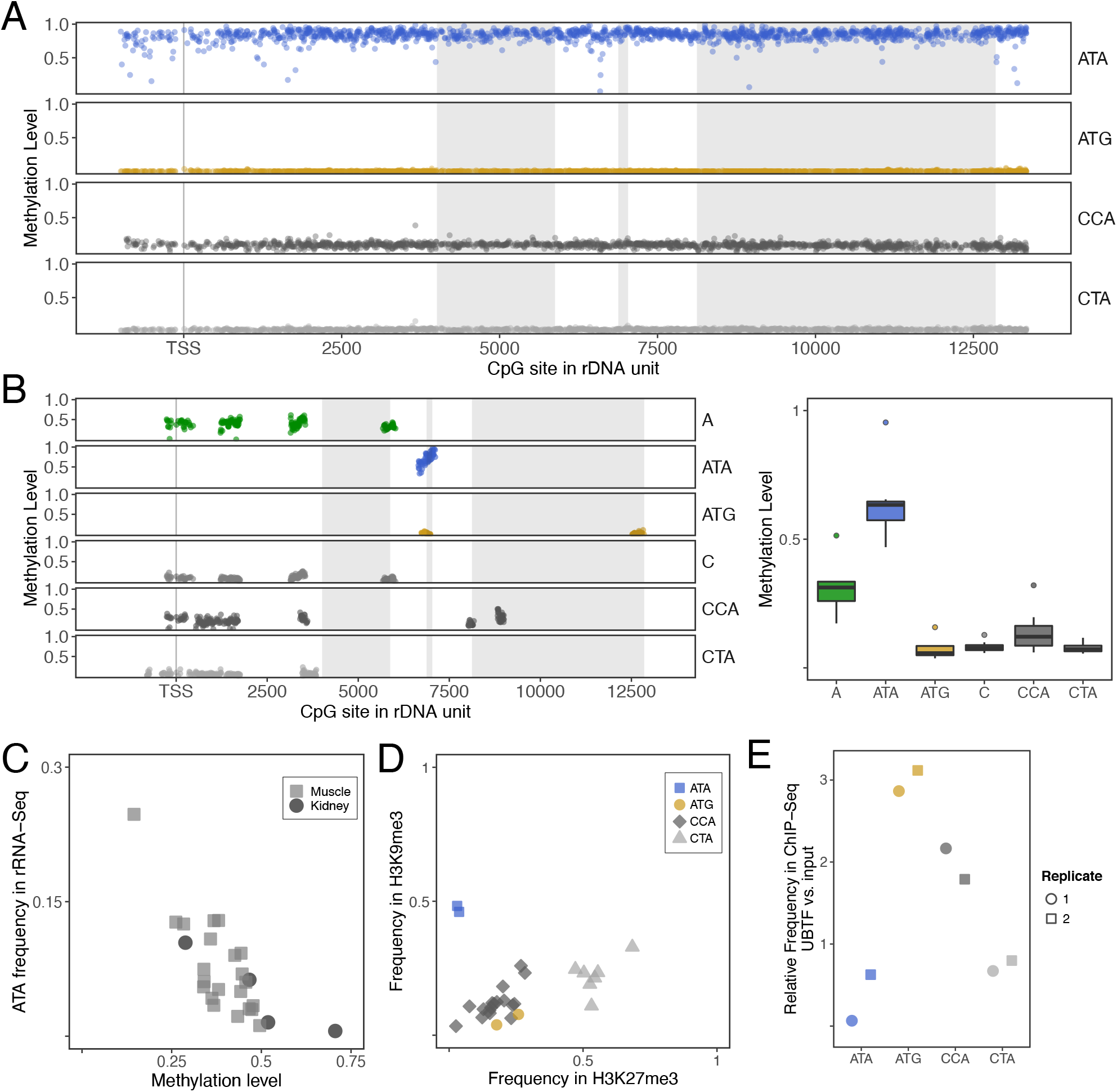
The functional genomic outcomes of C57BL/6J rDNA haplotypes. **(A)** Direct CpG methylation profiles of rDNA haplotypes from ultra-long read Nanopore MEF data, from −1000 bp upstream of the TSS up to the end of the 3’ ETS. **(B)** Example of rDNA haplotype-specific methylation analysis on C57BL/6J kidney whole genome bisulfite sequencing (WGBS) data (left panel) showing positional information for the CpG sites associated with each haplotype in a single mouse, and aggregate sperm Reduced Representation Bisulfite Sequencing (RRBS) data for seven mice (right panel). Note short-read technologies limit the range of positions that can be considered for each haplotype to the close neighbourhood of their uniquely-identifying SNVs. See also **Additional file 3: fig. S4**. **(C)** Relationship between A-haplogroup (“A” at −104) promoter methylation in C57BL/6J muscle (squares, n=22, p-value = 7.6×10^-6^, bis-PCR) and average coding + promoter methylation from kidney (circles, n=4, ATA-only WGBS) correlated with ATA-haplotype expression using rRNA-seq. **(D)** The ATA haplotype is enriched for H3K9me3 but depleted for H3K27me3 on CUT&Tag C57BL/6J kidney data (biological replicate - **Additional file 3: fig S5**). **(E)** The ATA haplotype is depleted for UBTF relative to other haplotypes in ChIP-Seq data from wild-type C57BL/6J B cells from Diesch *et al*., 2019^14^.

DNA methylation at the mouse rDNA promoter has a strong effect on rRNA expression^10^. If methylation is restricted largely to ATA haplotypes, then methylation observed at the rDNA promoter of ‘A’ haplogroup should derive from ATA haplotypes only, and correlate with the relative frequency of ATA in the rRNA. We performed ‘rRNA-seq’, which omits the rRNA depletion step in the standard mRNA-seq protocol (**Additional file 1**). Also, because of the sheer abundance of cellular rRNA, even internal transcribed spacer (ITS) RNA is readily detected (ITS2 contains the ATA-defining SNV at 6832). We observed a strong negative correlation between ‘A’ haplogroup promoter methylation levels and the relative frequency of A’s at position 6832 in muscle rRNA-seq (**Fig. 2C**; **Additional file 2: table S4**). In fact, since DNA methylation levels within any given coding unit are similar along the length of the unit, methylation even in the vicinity (+/- 200bp centred on the variant) of position 6832, in the rDNA should be correlated with the variant ratios observed at position 6832 in rRNA-seq, which is what we observe using 4 different kidney WGBS datasets (**Fig. 2C**). Next, we performed Cleavage Under Targets & Tagmentation (CUT&Tag - a recently developed method to profile histone modifications and chromatin proteins with improved signal to noise ratio^11^) analysis on kidney samples, finding that only ATA haplotypes display combined H3K9me3 enrichment/H3K27me3 depletion, consistent with previous reports of the relationship between DNA methylation and these histone modifications in other genomic regions^12^ (**Fig. 2D**; **Additional file 3: fig. S4**; **Additional file 2: table S5**). Finally, it is known that the Upstream Binding Transcriptional factor (UBTF) binding to unmethylated rDNA is required for rRNA expression^13^. Re-analysis of a previously published dataset of UBTF binding in B-cells of C57BL/6J mice^14^, showed that ATA is depleted for UBTF relative to the non-ATA haplotypes (**Fig. 2E**). Collectively, these analyses demonstrate key functional genomic features of ATA vs non-ATA rDNA haplotypes in a variety of tissue types.

### DNA methylation dynamics at rDNA are dichotomous and life-stage specific

Epigenetic silencing of rDNA is an integral component of the stress response in all eukaryotic cells. Recent studies using the C57BL/6J strain showed that pre-weaning exposure to nutritional stress, e.g., maternal protein restriction, obesogenic or high-fat diet, induces DNA hypermethylation of rDNA that persists into adulthood^6,7,15^. We re-analysed our previous RRBS data from C57BL/6J individuals exposed to maternal protein restriction in prior to weaning^6^, and found that the hypermethylation occurs specifically at the ATA variants (**Fig. 3A**; **Additional file 3: fig. S5**). In the GEO database, RRBS data is available for one other pure C57BL/6J mouse model of nutritional stress in which the offspring were exposed to a high fat diet during gestation+lactation, or during gestation+lactation+post-weaning^16^. In both experimental groups, relative to controls, rDNA hypermethylation is observed at ATA variants only (**Fig. 3A**). The early life nutritional stress models demonstrate that certain stimuli induce methylation and silencing of ATA rDNA units. But, can ATA units also lose methylation and re-activate? We re-analysed Dahlet *et al*.’s RRBS data of C57BL/6J CRISPR-based Dnmt1 knockout (Dnmt1 KO) 8.5dpc embryos^17^ and found that Dnmt1 KO embryos show virtually no methylation at any of the rDNA haplotypes (**Fig. 3B, left panel**; **Additional file 3: fig. S6**). In standard mRNA-seq protocols, rRNA is specifically removed. However, this depletion is never completely efficient and we find millions of rRNA reads in the Dahlet *et al.* mRNA-seq data (**Additional file 1**). Our own analysis of 22 matched C57BL/6J muscle mRNA-seq and rRNA-seq revealed an excellent correlation of rRNA variant frequencies (R ≥ 0.98 in all cases; **Additional file 3: fig. S7**). We then examined rRNA expression in Dahlet *et al.*’s mRNA-Seq data and found that ATA variants are expressed at considerably higher relative levels in the Dnmt1 KOs, proving that methylated ATA variants are not irreversibly silenced (**Fig. 3B**).

**Figure 3.**
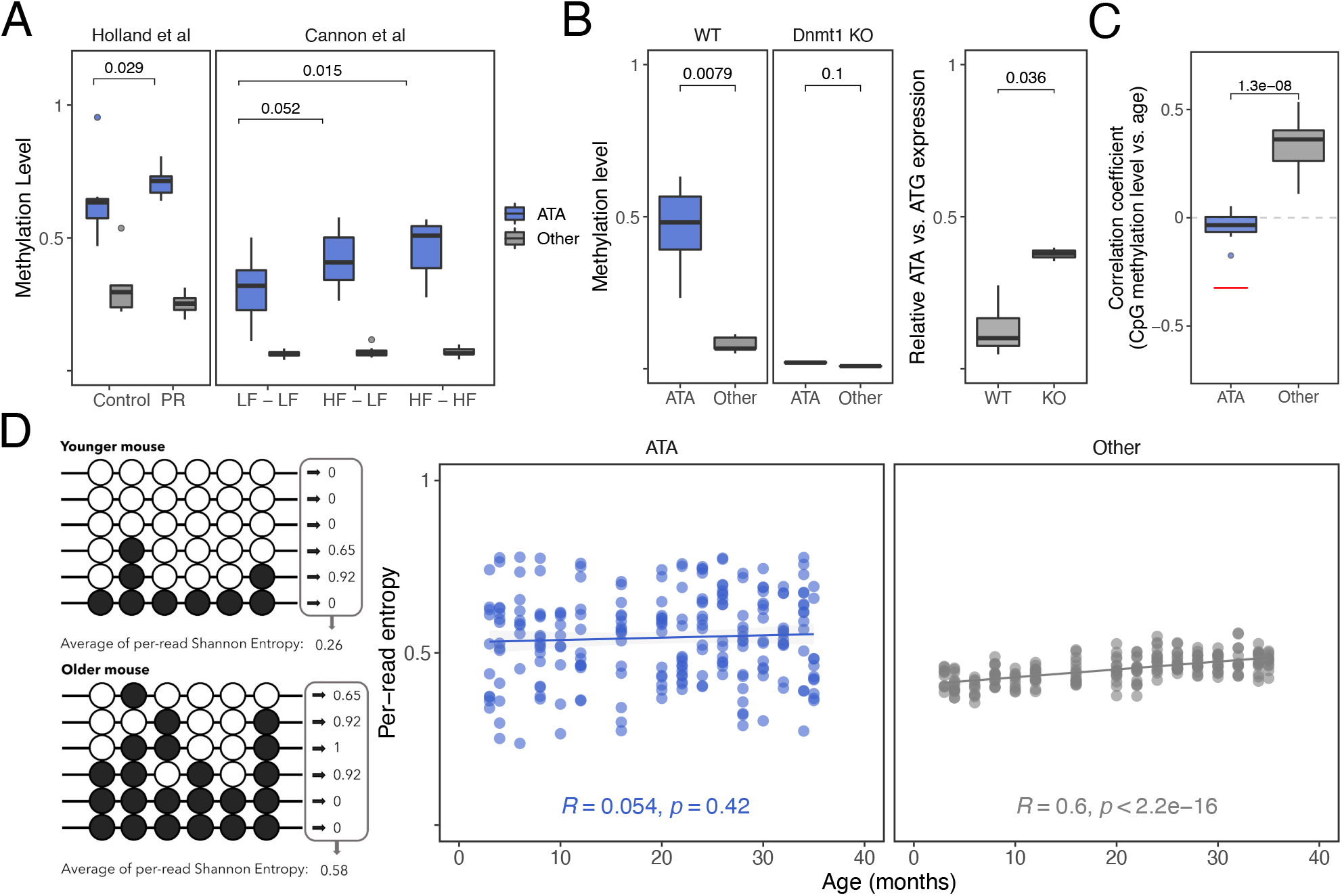
Epigenetic dynamics at C57BL/6J rDNA haplotypes are life stage specific. **(A)** Hypermethylation of the ATA haplotype (blue) is observed in different models of *in utero* stress paradigms; the other three haplotypes (grey) are combined into a single category since they show similar effects. Methylation levels for ATA and non-ATA haplotypes for all panels are obtained on the CpG sites associated with the SNVs at 6007 and 6832. PR, LF, and HF stand for Protein-restricted, Low-fat and High-fat diet, respectively. All animals from Holland *et al*.^6^ were fed control diet postweaning. The labels for Cannon *et al*.^16^ denote the pre-weaning, post-weaning diet combinations. **(B)** In Dnmt1 KO 8.5dpc embryos from Dahlet *et al*.^17^, the ATA haplotypes lose methylation in RRBS data (left panel) and display de-repression of ATA-haplotype rRNA expression (right panel). Both ATA and ATG expression are obtained from ITS2 variants to avoid differential rRNA depletion effects. **(C)** In RRBS data from Petkovich *et al*.^19^, ageing-associated DNA hypermethylation is observed only at non-ATA haplotypes. The red line indicates the expected correlation coefficient for ATA haplotypes if they lost methylation at the same rate as the non-ATA haplotypes gain methylation. In this study, in the 3-month old mice the average methylation at ATA was ~75% and at non-ATA haplotypes was ~25%. **(D)** Only non-ATA haplotypes display increasingly disordered DNA methylation profiles with age (right panel) in RRBS data from Petkovich *et al*.^19^. Disorder is estimated as the average of per-read Shannon entropy (left panel - example from two hypothetical mice).

Interestingly, previous studies show that a low protein diet in C57BL/6J mice post-weaning does not induce rDNA hypermethylation^7,18^. The study by Cannon *et al.* also includes a fourth group in which the mice were exposed to a high-fat diet post-weaning only, and in this group evidence for diet-induced epigenetic differences at rDNA is weaker^16^ (**Additional file 3: fig. S5**). To further explore the idea that the epigenetic state of ATA rDNA shows only limited dynamics in adulthood, we leveraged the RRBS dataset of Petkovich *et al*., 2016, representing 193 different C57BL/6J mice spanning an age range from 3-35 months^19^. Ageing-associated DNA methylation dynamics are observed genome-wide in a large range of mammalian species^20–22^, including at rDNA in humans and rodents^23–26^. In the Petkovich *et al.* dataset, ATA showed no directional change with age (**Fig. 3C**; **Additional file 3: fig. S8, S9**). However, methylation at non-ATA haplotypes, and in particular CCA and CTA, display a positive correlation with ageing (**Fig. 3C**). By leveraging single-molecule level data, we found that this was primarily driven by an increase in DNA methylation entropy within individual DNA molecules at non-ATA haplotypes (**Fig. 3D**; **Additional file 3: fig. S10**, **S11**)^27–29^. Therefore, epigenetic dynamics at rDNA haplotypes are life-stage specific and dichotomous: the ATA haplotype displays environmentally-induced epigenetic dynamics during early development, but is less susceptible to further perturbations in later life. On the other hand, non-ATA haplotypes entropically accumulate methylation during ageing.

### rDNA epialleles are present in other mouse strains and human

We then studied 5 additional inbred mouse strains (**Fig. 4A**). For each strain, we generated kidney WGS, WGBS, rRNA-seq, and droplet digital PCR (ddPCR) data from each of 6 different adult males (117 different datasets, as 3 were discarded post-QC) (**Additional file 2: table S3**). We confirmed that for any given sample, >82% of the rDNA coding unit SNVs called in the WGS data are also found in the WGBS data (**Additional file 2: table S6-10**; **Additional file 3: fig. S12**). We then asked which of these SNVs are associated with allelic methylation differences, i.e., are ‘epivariants’ (Wilcoxon rank sum test, FDR < 0.01), strain-specific and/or common across multiple strains (**Fig. 4A**; **Additional file 3: fig. S13**; **Additional file 2: table S11**). Analysis of matched rRNA-seq data showed that epivariant-associated methylation differences throughout the coding unit impact variant frequencies in the rRNA, with the exceptions being 129S1/SvImJ and C3H/HeJ (**Fig. 4B**; **Additional file 3: fig. S14**; p-value=3.464×10^-6^, paired Wilcoxon rank sum test on difference of correlation coefficients). We noted that these two strains also showed the lowest levels of rDNA methylation (**Fig. 4A**). Given the known positive correlation between increasing total rDNA copy number (CN) and epigenetic silencing in lower organisms^30^, we considered the possibility of that CN might account for strain-specific differences in total rDNA methylation levels. Therefore, we calculated CN from three independent datasets - WGS, WGBS and ddPCR - for each individual mouse, as CN measurements in mammals are known to be technically challenging^31^ (see **Additional file 1** for CN calculations; **Additional file 3: fig. S15**). Indeed, we found a significant positive correlation between total rDNA CN and DNA methylation across the 6 different inbred strains (**Fig. 4C**; **Additional file 3: fig. S15**).

**Figure 4.**
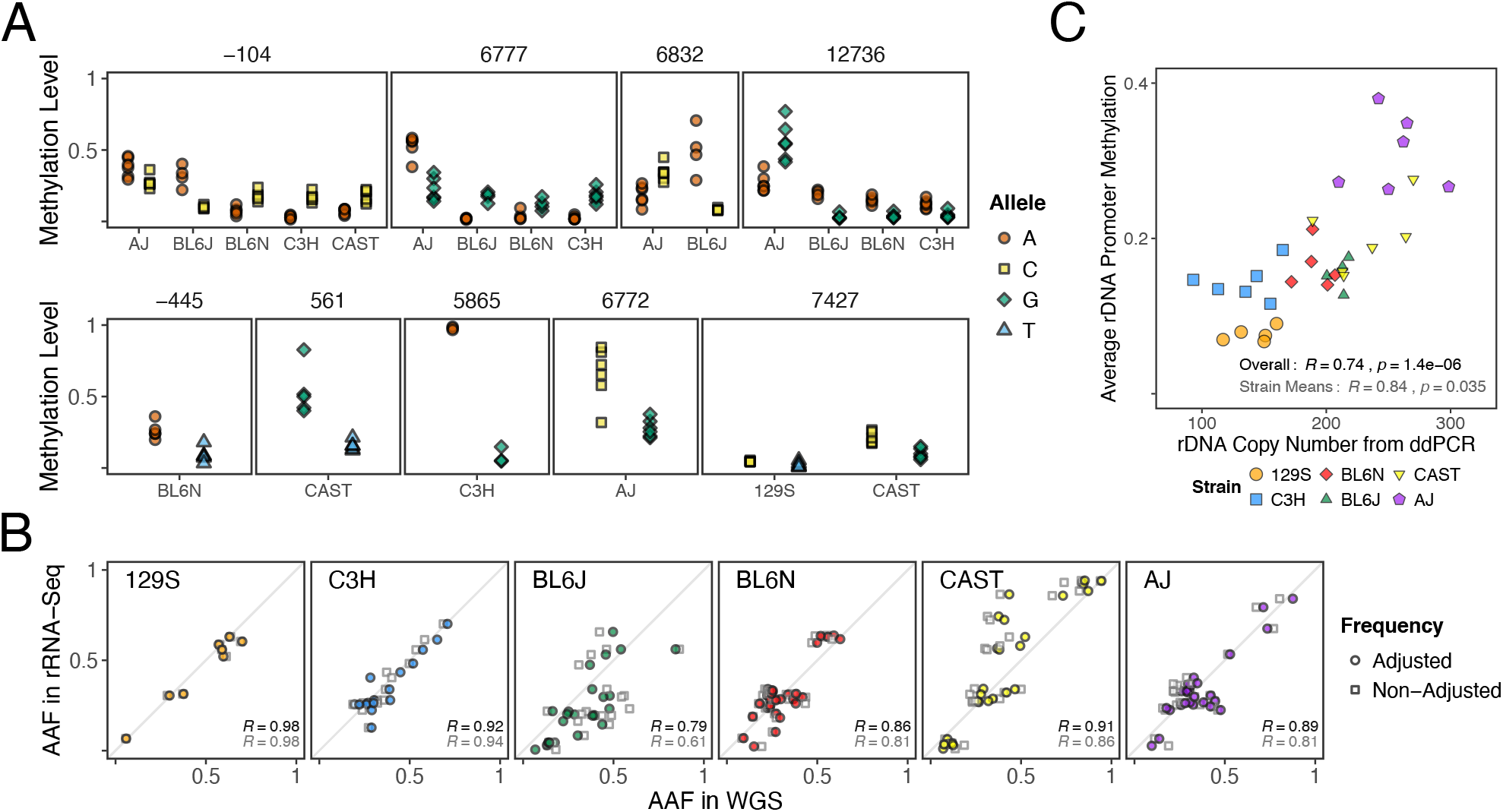
The epiallelic nature of rDNA is a feature of other mouse inbred strains. **(A)** Allelespecific methylation levels at selected rDNA epivariants from kidney of 5 inbred strains besides C57BL/6J (‘BL6J’): 129S1/SvImJ (‘129S’), C3H/HeJ (‘C3H’), C57BL/6N (‘BL6N’), CAST/EiJ (‘CAST’) and A/J (‘AJ’) (see **Additional file 3: fig. S13** for a comprehensive display of all identified epivariants). The top panel shows positions associated with C57BL/6J haplotypes. Notably, the methylation differences at these positions are not always directionally consistent across different strains where the variants are conserved. The bottom panel depicts strain-specific variants; note 7427 is variable in 2 strains, but with different alternative nucleotides (T in 129S1/SvImJ and G in CAST/EiJ). **(B)** Comparison of alternative allele frequencies (AAF) at the epivariant positions between rDNA (WGS) and rRNA for one representative mouse of each strain (see **Additional file 3: fig S14**). If only unmethylated units are expressed, then adjusting the AAF ratios in the WGS by ignoring methylated copies in the ratio calculations should improve the correlation between variant frequencies (i.e. unmethylated correlated with expressed). This is observed in 4/6 strains. **(C)** Total rDNA copy number from ddPCR correlated with methylation level from WGBS across the different mouse strains. Correlation coefficients displayed use all mice as data points (black) or a single average point per strain (grey). The analysis was repeated using the highly conserved 18S subunit only to exclude any possibility that the results are due to strain-specific differences in mapping efficiency (see **Additional file 3: fig. S15**).

To ask if human rDNA displays similar genetic-epigenetic relationships, we generated WGBS and rRNA-seq data for 48 different human lymphoblastoid cell lines (LCLs) derived from ‘Gambian in Western Division - Mandinka’ adult individuals that were sequenced in the 1000 genomes project^32^ (**Additional file 2: table S12**). We first confirmed that for any given sample, >95% of the rDNA coding region SNVs called in the published WGS data were also called in our WGBS data (**Additional file 3: table S12**). Even in this relatively small number of samples for an outbred population, we identified two epivariants at FDR<0.01 (**Fig. 5A**). DNA methylation in the vicinity of these sites (−413 located in the promoter and 7980 located in the 28S) was positively associated with total rDNA CN (**Fig. 5B**). We then analysed the 7980 variant in rRNA-seq datasets for these samples and found DNA methylation in the vicinity of 28S negatively associated with variant representation in rRNA (**Fig. 5C**). Finally, we re-analysed published ultra-long read Nanopore data from a GWD sample (**Additional file 1**). Interestingly, in contrast to the mouse rDNA, we did not find extensive evidence for genetic haplotypic structure within a unit. However, comparison of adjacent units revealed strong epigenetic relatedness like in the mouse, and also some evidence that at least the 7980 position shows genetic similarity between neighbouring units (**Fig. 5D**).

**Figure 5.**
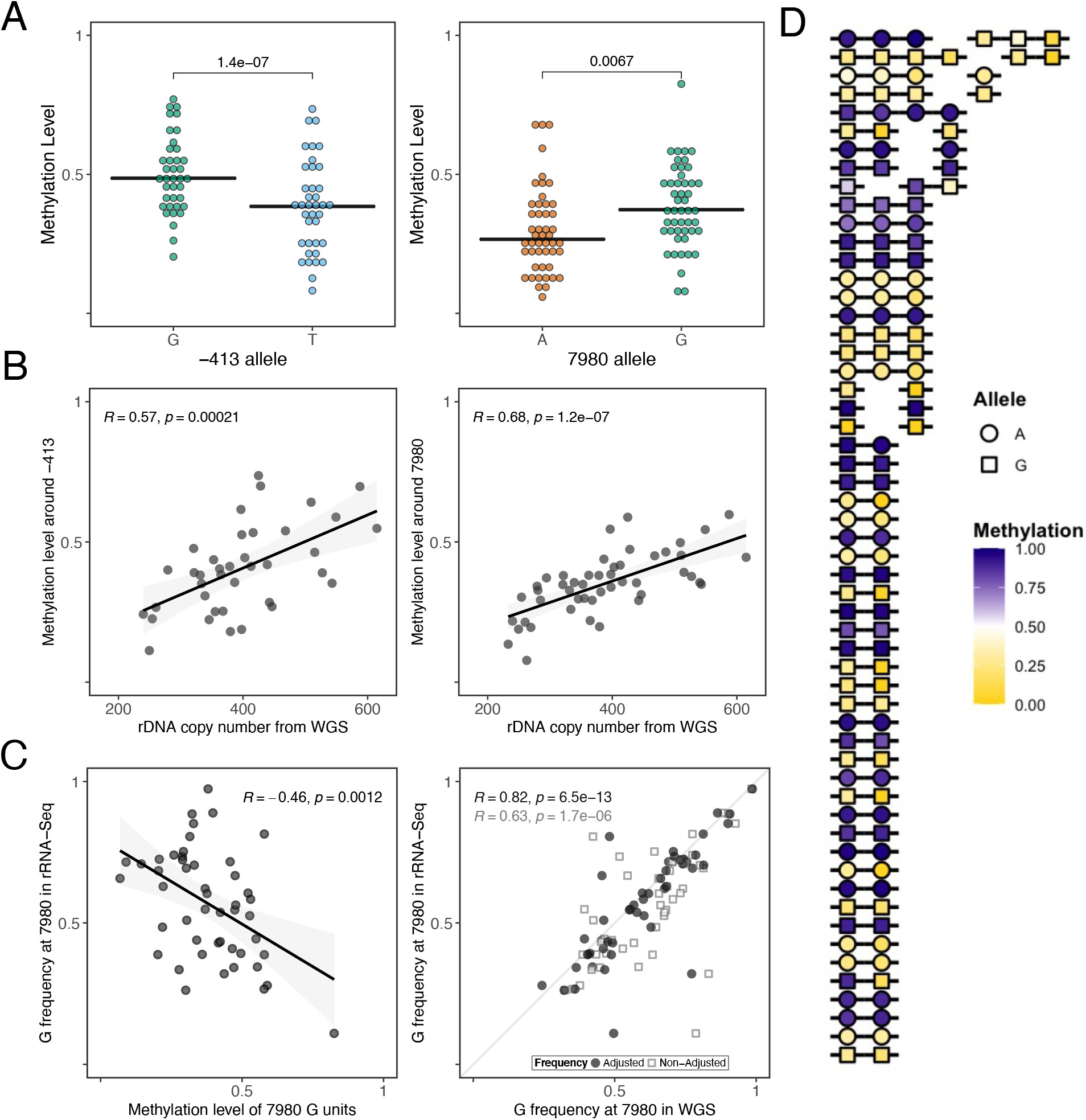
Human Mandinka samples display rDNA epivariation. **(A)** Analysis of human Mandinka LCL WGBS samples shows allele-specific methylation levels (FDR < 0.01) at positions - 413 and 7980 of the KY962518.1 rDNA reference. **(B)** Average methylation levels around −413 and 7980 are correlated with rDNA copy number estimated from WGS. **(C)** Allele-specific methylation affects rRNA expression. Methylation levels around position 7980 are inversely correlated with allele-specific frequency in rRNA-seq (left panel), and correlation between DNA (WGS) and rRNA allele frequencies improves when accounting for methylation at position 7980 (right panel). **(D)** Analysis of ultra-long read Nanopore data from human sample HG02723 suggests that both allele at position 7980 (shape) and average methylation level across the rDNA coding unit (fill colour) tend to be consistent along single molecules, with some exceptions. Each row represents an individual read; gaps separate units that were reported on split alignments.

## Discussion

Here we have shown that the genetic identity of a rDNA unit has a strong probabilistic influence on the homeostatic epigenetic state. The CN data suggests that the emergence of rDNA epialleles could be linked to the need of the genome to silence extra copies of rDNA. However, the silencing is not absolute, reminiscent of the epigenetic metastability associated with some mammalian retroelements^33,34^. Could the loss/gain of specific rDNA variants underlie the CN differences? It has been shown that the number of different chromosomes harbouring rDNA clusters can vary from 3-5 amongst different inbred mouse strains^35^. The C57BL/6J genome contains rDNA clusters on 4 separate chromosomes^35^, leading to the possibility that each chromosome harbours a different variant given that we find 4 different rDNA haplotypes in the C57BL/6J strain. Interestingly, the C3H strain has only 3 rDNA clusters^35^, and shows very few DNA methylation-based rDNA epialleles.

But could non-rDNA methylation based epialleles exist in the mouse? We performed CUT&Tag analysis for H3K9me3 and H3K27me3 in the 129S1/SvImJ strain which has very few DNA methylation-based epivariants in rDNA, and even these are of very small effect size. However, we did not find any patterns/clusters of the histone modifications (unlike what is observed for C57BL/6J), suggesting that there are no preferentially epigenetically silenced genetic variants in the absence of DNA methylation, at least in 129S1/SvImJ mice (**Additional file 3: fig. S16**). Our 129S1/SvImJ data also suggests that in some mouse strains, DNA methylation plays little, if any, role in silencing of rDNA copies.

For human rDNA, we found limited evidence of genetic haplotypes within a unit, or genetic similarity of adjacent units, consistent with a recent study^36^. It is previously been noted that inter-chromosomal rDNA recombination is greater in human than in the mouse^37^, which would be consistent with our findings. However, there was clear evidence for epiallelic effects within a unit, albeit not as marked as for the mouse, and also epigenetic states of neighbouring human rDNA units are similar. This raises the intriguing possibility that the entire cluster of rDNA units on any given chromosome share a similar epigenetic identity, and this property is conserved amongst mammalian genomes. In the future, it will be interesting to analyse other mammalian genomes to elucidate what determines higher order genetic and/or epigenetic patterns of mammalian rDNAs.

With regards to the molecular consequences of rDNA (epi)genetic variation in models of nutritional stress and ageing, a tantalizing possibility is the production of variant ribosomes that have subtle but measurable effects on translational output, in line with the ribosome filter hypothesis^1,4^. It has been shown that endogenous rRNA sequence variation regulates stress response gene expression in bacteria^38,39^. Previous studies have shown that human and mouse coding subunit rDNA genetic variation is located in rRNA regions with known roles in translation^1^. An initial Polysome-seq analysis shows that the ratio of 28S rRNA variants incorporated into ribosomes is dependent on the ratio at which they are expressed (**Additional file 3: fig. S17**). In eukaryotes, there are examples of variation in either ribosomal proteins or rRNA modifications influencing preferential translation of some mRNAs^1^, yet the contribution of rRNA sequence variation has remained largely unexplored and will require large-scale translatome analyses in the future.

## Conclusions

Here we have shown that genetic variation at mouse and human ribosomal DNA influences epigenetic states and associated transcriptional outcomes. Currently, this source of genetic variation is largely overlooked in large scale studies and thus it is possible that rDNA-associated genetic variation underlies some apparently ‘epigenetic’ phenomena^40^ and/or contributes, *in trans*, to the ‘missing heritability’ in some human phenotypes and diseases.

## Methods

### Mouse embryonic fibroblasts

were made from a 13.5 dpc male C57BL/6J embryo following the protocol of the Jacks Lab (available online at http://web.mit.edu/jacks-lab/protocols/Making_MEFs_tables.html). Immortalisation was done using the PA317 (λtsA58U19-neo) (G418 0.75mg/ml) viral vector.

### Ultra-long Nanopore sequencing

was performed using the protocol of Jain *et al*.^9^.

### Short read sequencing libraries

were generated using the following kits and according to the manufacturers protocols: (i) Whole genome sequencing (WGS) - NEBNext^®^ Ultra™ II DNA Library Prep Kit for Illumina, NEB, USA; (ii) Whole genome bisulfite sequencing (WGBS) - Accel-NGS^®^ Methyl-Seq DNA Library Kit, Swift BioSciences, USA; RNA-seq - NEBNext^®^ Ultra™ II Directional RNA Library Prep Kit for Illumina, NEB, USA. rRNA-seq libraries were made like the RNA-seq but without any depletion. All sequencing was performed on an Illumina Novosaq6000 by Novogene, Cambridge, UK. For WGS and WGBS, we generated data in the range of ~15-25X genome-wide coverage on average, corresponding to ~350-1350X at the rDNA. We have previously shown that this level of sequence coverage at rDNA results in data that is extremely well correlated with targeted PCR based approaches (in which the coverage is not limited)^6,7^. The methylation values for muscle in **Fig. 2C** were derived from multiplex bisulfite PCR data generated in Reference 7. Bisulfite PCR Sequencing (Bis-PCR-seq) was performed on DNA from muscle, as in Reference 7. DNA was bisulfite converted using the EZ-96 DNA Methylation™ Kit (Zymo, Cat. D5003). Targeted amplification was performed using the FastStart High Fidelity PCR System, dNTPack (Sigma-Aldrich, cat. 4738284001) in the 48.48 layout on the Fluidigm C1 system (Fluidigm, USA), a microfluidics platform. Library preparation was performed using the same kit including 4 ul of Access Array BArcode Library Primer and 1 ul of PCR product diluted 1:100. Libraries were sequenced with Illumina NextSeq (75 bp, single-end).

### Droplet digital PCR

was used to measure total rDNA copy number across different strains using mouse kidney tissue. Probes and primers were designed against the genomic mouse 18S rDNA sequence (GenBank: BK000964.3, positions 4008-5877). The 18S targeting probe was attached to the 5’ fluorescent dye, FAM, and a 3’ nonfluorescent quencher, NFQ. The mouse transferrin receptor, Tfrc, was used as a single copy reference, and was targeted by a pre-designed assay with a HEX fluorescent dye and an Iowa Black quencher (BioRad, Assay ID: dMmuCNS420644255). The restriction enzyme, AluI (NEB, Cat. RO137S) was used to separate each rDNA gene copy to minimise the number of copies per droplet. AluI was diluted with 1X CutSmart Buffer to give a concentration of 1U/ul. 50ng of DNA was used for every AluI digestion. 20 units of AluI for every 1ug of DNA was used and the reaction was incubated at 37 °C for 1 hour. 1ng of digested DNA was then added to each sample well. The ddPCR reactions were prepared in duplicate in a ddPCR™ 96-well plate (Bio-Rad, Cat. 12001925). The components of the reaction mix per reaction were 11ul iTaq Universal Probe Supermix (BioRad, Cat. 1725130), 0.275ul FAM, 1.926ul HEX and an 8.8ul solution of water and DNA. The automated procedure of ddPCR was carried out using the QX200™ AutoDG™ ddPCR system from Bio-Rad. The automated system involved a droplet generator machine (Bio-Rad, Cat. 1864101) DG32™ Automated Droplet Generator Cartridges (Bio-Rad, Cat. 1864108), Automated Droplet Generation Oil for Probes (Bio-Rad, Cat. 1864110) and Pipet Tips for the AutoDG™ System (Bio-Rad, Cat. 1864120). The new plate with droplets was sealed with a pierceable PCR foil at 180 °C for 5 seconds using PX1 PCR Plate Sealer (Bio-Rad, Cat. 1814000). The PCR reaction was then carried out using the C1000 Touch™ Thermal Cycler (Bio-Rad, Cat. 1851197). PCR settings were: 10 minutes 95°C for initial denaturation, cycle of 40 repetitions, 94 °C for 30 seconds, annealing temperature: 57 °C and lid temperature: 105 °C. After PCR, the plate was processed in a plate reader and the data was generated with Quantasoft Software.

### CUT&Tag-seq

was performed according to the protocol of Kaya-Okur *et al*.^11^ with modifications to tissue processing as described below. Experiments were performed in duplicate from two independent mouse kidney tissues in parallel to minimize technical variation. To adapt CUT&Tag for mouse tissue sections, flash frozen mouse liver tissues (approximately 3-4mm size) were manually homogenised with tight homogenisers in wash buffer (20mM HEPES pH7.5, 150 mM NaCl, 0.1% BSA, 0.5mM Spermidine and cOmplete EDTA free protease inhibitor tablet) into a homogenous suspension of intact cells. Cells were transferred to 1.5 ml low DNA binding tubes (Eppendorf), and solutions were exchanged on a magnetic stand (DynaMag-2, ThermoFisher scientific). Cells were pelleted by centrifugation for 3 min 600 x g at room temperature and resuspended in 500ul of ice-cold NE1 buffer (20 mM HEPES-KOH pH 7.9, 10 mM KCl, 0.5 mM spermidine, 1% Triton X-100, and 20 % glycerol and cOmplete EDTA free protease inhibitor tablet) and let it sit for 10 min on ice. Nuclei were pelleted by centrifugation for 4 min 1300 x g at 4°C and resuspended in 500ul of wash buffer and remove the wash buffer by placing the tubes on a magnet stand to clear and withdraw the liquid. Resuspended in 1.0 mL wash buffer and held on ice until beads are ready. 10 ul of BioMag Plus Concanavalin-A-conjugated magnetic beads (ConA beads, Polysciences, Inc) in binding buffer (20 mM HEPES-KOH pH 7.9, 10 mM KCl, 1mM CaCl_2_ and 1mM MnCl_2_) was added to each tube containing cells and rotated on an end-to-end rotator for 10 minutes. After a quick spin to remove liquid from the cap, tubes were placed on a magnet stand to clear and withdraw the liquid, and 800ul of antibody buffer containing 1ul of primary antibodies (normal rabbit IgG, Santa Cruz Cat no sc-2027, H3K27me3 (Millipore, Catalogue number 07-449, Lot DAM1703508 and H3K9me3 (Diagenode, Catalogue number C15410193, Lot A.0219P) was added and incubated at 4°C overnight in a nutator. Secondary antibodies (guinea pig α-rabbit antibody, Antibodies online cat. no. ABIN101961) was added 1:100 in Dig-wash buffer (5% digitonin in wash buffer) and squirt in 100 μL per sample while gently vortexing to allow the solution to dislodge the beads from the sides and incubated for 60 min on a nutator. Washed unbound antibodies in 1 ml of Dig-wash buffer for a total of three times. 100 ul of (1:250 diluted) protein-A-Tn5 loaded with adapters (kind gift from Steven Henikoff lab) in dig-300 buffer (20 mM HEPES pH 7.5, 300 mM NaCl, 0.5 mM spermidine with Roche cOmplete EDTA free protease inhibitor) placed on nutator for 1 hour. Washed three times in 1 ml of Dig-300 buffer to remove unbound pA-Tn5. 300 μL Tagmentation buffer (Dig-300 buffer + 5 mM MgCl_2_) was added while gently vortexing and incubated at 37°C for 1hr on an incubator. Tagmentation was stopped by adding 10 μL 0.5M EDTA, 3 μL 10% SDS and 2.5 μL 20 mg/mL Proteinase K to each sample. Mixed by full speed vortexing for ~2 seconds and incubate 1 hr 55°C to digest. DNA was purified by phenol:chloroform extraction using phase lock tubes followed by ethanol precipitation. Libraries were prepared using NEBNext HiFi 2x PCR Master mix (Cat number M0541S) with 72°C gap filling step followed by 13 cycles of PCR with 10 second combined annealing and extension for enrichment of short DNA fragments.

### Preparation of RNAs from polysome fractions

Mouse liver tissue extracts were prepared as previously reported^41^, using polysome extraction buffer (20 mM Hepes-NaOH (pH 7.4), 130 mM NaCl, 10 mM MgCl2, 1% CHAPS, 0.2 mg/ml heparin, 5% glycerol, 2.5 mM DTT, 50 μg/ml cycloheximide, 20 U SUPERase In RNase inhibitor, cOmplete EDTA-free Protease inhibitor). Ribo Mega-SEC run was carried as previously reported^42^, by the flow rate of 0.4 ml/min, and polysome fraction (1.2 ml) was collected from 16.5 min to 19.5 min. RNAs in the polysome fraction were extracted by TRIzol LS reagent (SIGMA) and precipitated with isopropanol containing glycogen. Precipitated RNAs were purified further by LiCl precipitation.

### Mouse strains

used in this study were ordered from Charles River, UK. All mice were 6 weeks of age when delivered, were allowed to acclimatize for 2 weeks, after which they were killed by Carbon dioxide. The mice used in **Fig. 2C**, and **Additional file 3: fig. S7** and **S17** are from Reference 7. DNA and RNA from tissues was extracted using Qiagen kits and following the manufacturers protocols. Aside from the C57BL/6J mouse strain, the other mouse inbred strains were selected to span a range of genetic and copy number variation in the rDNA. Strain-specific copy number estimates from Parks *et al*.^2^, and WGS sequencing data from the Mouse Genomes Project were obtained for 11 different strains: 129S1/SvImJ, A/J, BALB/cJ, C3H/HeJ, C57BL/10J, C57BL/6N, CAST/EiJ, CBA/J, DBA/2J, FVB/NJ and MOLF/EiJ^43^. Sequencing reads were processed as indicated below to obtain rDNA SNVs for each strain (see “Short read data processing” in **Additional file 1**). Allele frequencies at all such SNV positions were then used to cluster strains with `R`’s Ward hierarchical clustering method (**Additional file 3: fig. S18**). Alongside copy number estimates, these clusters served as the main support for selecting 129S1/SvImJ, A/J, C3H/HeJ, C57BL/6N, and CAST/EiJ as additional strains to consider in the current study.

### Human lymphoblastoid cell lines (LCLs)

of the Gambian in Western Division - Mandinka (GWD) population obtained from the Coriell Institute (New Jersey, United States), were used for all human experiments. Cell lines were seeded at a density of ~200,000 cells/ml in RPMI 1640 + GlutaMAX (Gibco; 61870-010) supplemented with 15% fetal bovine serum (Gibco; 10270-106) and a 1% penicillin and streptomycin mix (Gibco; 15140-122). All cell cultures were kept in 37°C incubators under 5% carbon dioxide conditions. For all down-stream experimental analyses, cell lines were pelleted, washed twice through with phosphate buffered saline (Sigma; P4417) and stored at −80C. Whole genome sequencing data from the 1000 genomes project was obtained for the 119 adult samples from the Gambian Mandinka population (GWD) with cell lines available from the Coriell Institute for Medical Research (all except HG02756). Sequencing reads were processed as indicated below to estimate both rDNA variation and copy number (see “Short-read data processing” and “Total rDNA copy number analysis” in **Additional file 1**). 24 male and 24 female samples were then selected to maximise the range of rDNA variation and copy number in further analyses (**Additional file 2: table S12**).

### Data Analysis

Unless explicitly stated, all analyses were conducted using in-house scripts implemented in `R` version `4.0.2.` An extended description of the data analysis methods in this study can be found in the **Additional file 1**, including detailed explanations of the rDNA reference sequences, command line parameters, and mathematical formulae employed.

Short-read sequencing data were first analysed using `fastqc` version `0.11.9` to identify potential issues, and successful libraries were then trimmed for both base quality and adaptor removal using `trimgalore` version `0.6.5.` Alignments to the reference sequences were performed using `bowtie2` version `2.4.1` for non-bisulfite-converted DNA data (WGS, ChIP-Seq, and CUT&Tag), `bismark` for bisulfite-converted data (version 0.7.12 for Bis-PCR-seq and version `0.22.1` for WGBS, RRBS) with underlying `bowtie2`, and `STAR` version `2.7.0f` for RNA data. Alignment output files were then sorted, indexed and filtered to retain only reads aligned to the appropriate rDNA unit reference using `samtools` version `1.10.` SNVs were called on all non-bisulfite-converted datasets using `lofreq` version `2.1.5.` For WGBS and RRBS data, CpG methylation estimates were obtained from the `bismark` alignments and then fed to `blink`, an in-house `python` tool for estimating allele-specific methylation and SNV frequencies, described more in detail in **Additional file 1** (see “`blink`: Allele-specific methylation and frequency from bisulfite-converted data”). `blink` was employed to obtain allele-specific methylation values from both mouse kidney and human LCL data (see “Analysis of mouse strains kidney data” and “Analysis of human LCL data”, respectively, in **Additional file 1**).

Total rDNA copy number estimates from ddPCR were obtained using the Quantasoft software in default mode. Estimates from short-read sequencing data (WGS and WGBS), on the other hand, were obtained following the procedure Gibbons *et al*^44^ suggest (see “Total rDNA copy number analysis” in **Additional file 1**).

Ultra-long read Nanopore libraries sequenced from MEFs were initially basecalled using `albacore` version `2.4.0` and aligned to the 18S and 28S regions from the BK000964.3 mouse rDNA reference using the `minimap2 python` interface `mappy`. The raw signal `fast5` files for reads mapping to such regions were later re-basecalled using `guppy` version `4.2.2` and mapped to a whole-genome plus rDNA mouse reference using `minimap2` version `2.17.` Genetic variation and CpG methylation were estimated from the rDNA reads using `megalodon`, whose output files were further processed as described in **Additional file 1** (see “MEF ultra-long read Nanopore data processing and analysis”) to obtain putative rDNA haplotypes and their corresponding methylation levels. Haplotype-specific alleles indentified from the MEFs were then employed to assign short Illumina reads to haplotypes (see “Haplotype-specific analysis of C57BL/6J kidney and muscle data” and “Processing and analysis of C57BL/6J publicly-available datasets” in **Additional file 1**).

Publicly available MinION ultra-long read Nanopore data for human sample HG02723 was obtained from Circulomics (https://www.circulomics.com/datasets). Raw `fast5` files were basecalled using `guppy` version `4.2.2` and aligned to the human whole-genome plus rDNA reference sequence with `minimap2`. Reads mapping to the rDNA were considered for further analysis with `megalodon` version `2.2.9`, using a reference sequence artificially expanded to include 20 back-to-back rDNA units. Short-read SNVs were obtained by first reconstructing with `samtools fastq` the raw sequencing reads from the CRAM file available from the 1000 genomes project, and then trimming, aligning and variant calling as previously described. These SNVs were then used as input in `megalodon` to obtain per-read variant and CpG methylation call, which were subsequently processed as described in **Additional file 1** (see “Human ultra-long read Nanopore data processing and analysis”) to establish potentially haplotypic structures.

## Declarations

## Acknowledgments

We thank Profs Sue Ozanne and Oliver Rando for feedback on the work, Dr Miguel Branco for the H3K9me3 Ab, and Dr Pankaj Dubey for help in CUT&Tag-seq library preparation. We also thank The Bart’s and the London Genome Centre for sequencing and the assistance of the ITS Research team at Queen Mary University of London.

## Author contributions

Conceptualization: VKR; Methodology: AFD, RAES, HY, PPL, AH, SJM, MMP, MLH, ML, VKR, Software: FRA; Formal Analysis: FRA; Investigation: SY, AFD, RAES, HY, PPL, ZA, PD, NH, VAM, AB, MMP, MLH, ML, VKR; Resources: ML, MLH, VKR; Data Curation: FRA, SJM; Writing - original draft: FRA, VKR; Review & editing: FRA, SY, AFD, RAES, HY, PPL, PD, SJM, MMP, AH, MLH, ML, VKR; Visualization: FRA, VKR; Supervision: ML, MLH, VKR; Project administration: VKR; Funding acquisition: MMP, MLH, ML, VKR

## Ethical approval

No ethical approval was required for this study.

## Availability of Data and Materials

All data generated for this study is available from BioProject ID: PRJNA733656.

## Additional files

**Additional file 1:** Extended data analysis methods

**Additional file 2:** Supplementary tables S1-S14

**Additional file 3:** Supplementary figures S1-S20

