## Additional File 1 - Extended Methods for "Genetic variation at mouse and human ribosomal DNA influences associated epigenetic states"

**Extended data analysis methods**

### **Reference sequences.** In order to reduce loss of coverage and spurious alignments, reference sequences employed in this study were created as follows. From the publicly available annotations for mouse (Genbank accession BK000964.3) and human (Genbank accession KY962518.1) rDNA unit references, the IGS repetitive element closest to the 3’ end of the unit was identified. The midpoint of such repetitive elements – located 3008 and 2120 base pairs upstream of the TSS, respectively – served as breakpoint to construct a so-called “looped” reference unit: the sequence from the breakpoint to the end of the original unit was prepended to the sequence from the TSS up to (but not including) the breakpoint. This avoids sequencing reads spanning both upstream and downstream of the TSS being discarded due to split alignments, improving read coverage in the region.

Sequencing reads originating from genomic regions other than the rDNA may be spuriously mapped to the rDNA, particularly on bisulfite-converted data. To minimise this risk, the “looped” rDNA unit above was appended to a whole genome assembly (MM10 for mouse and Hg38 for human) with identified rDNA pseudocopies masked, including the entire unit present in an unplaced Hg38 contig (Genbank accession GL000220.1; see Reference 45). **Table S14** from **Additional file 2** lists the genomic coordinates for all masked regions in both mouse and human assemblies. These joint “masked” whole genome plus “looped” rDNA reference sequences were then used for subsequent short- and long-read analyses.

Gene annotations were obtained from Gencode – vM25 for mouse and v34 for human – in gtf format, and augmented with entries for rDNA components. These gene annotations were then attached to the reference sequences above using STAR --runMode genomeGenerate with parameter --sjdbOverhang 149, creating reference sequences suitable for RNA-seq alignments.

Exome references were also obtained and adapted, following the procedure Gibbons *et al*. describe^44^. Exome sequences and corresponding annotations – labelled 10090 for mouse and 9606 for human – were downloaded from the EMBL/EBI online repository. Sequences from the sex chromosomes and smaller than 300 base pairs were removed. Significantly similar sequences, as reported by blastn version 2.7.1+ in --ungapped mode, were also filtered out, leaving a total of 14,148 sequences in the mouse reference and 12,898 in the human reference.

**Short-read data processing**. Short-read sequencing data were first analysed using fastqc version 0.11.9 to identify potential issues, and successful libraries were then trimmed for both base quality and adaptor removal using trimgalore version 0.6.5, with the –-paired and ‑‑rrbs options enabled when appropriate. For the WGBS mouse kidney and human datasets, the parameters ‑‑clip_r1 10 and ‑‑clip_r2 20 were also included as recommended for their library preparation kits. All remaining parameters were set to their default values. Trimmed reads were then further analysed using fastqc.

Alignments to the reference sequences were performed using bowtie2 version 2.4.1 for non-bisulfite-converted DNA data (WGS, ChIP-Seq, and CUT&Tag), bismark for bisulfite-converted data (version 0.7.12 for Bis-PCR-seq and version 0.22.1 for WGBS, RRBS) with underlying bowtie2, and STAR version 2.7.0f for RNA data. All bowtie2 and bismark alignments were conducted using default parameters. Bisulfite-converted datasets were aligned to reference sequences converted *in‑silico* using bismark_genome_preparation. In STAR, the options -‑outSAMstrandField intronMotif and --quantMode GeneCounts were requested. Alignment output files were then sorted, indexed and filtered to retain only reads aligned to the appropriate rDNA unit reference using samtools version 1.10.

SNVs were called on all non-bisulfite-converted datasets using lofreq version 2.1.5, which provides allele frequency estimates without assuming an underlying diploid genome. rDNA alignments were first quality-annotated using lofreq alnqual and then fed to lofreq call, limiting the region of interest up to the end of the 3’ ETS, in two different modes: (i) exhaustive, with parameters --call-indels -a 1 -b 1 -B ‑-no-default-filter -‑use-orphan, and (ii) default, with only --call-indels enabled. The output of the latter mode was then further processed using lofreq filter --only-snvs to produce both “high confidence” (parameters -v 250 -a 0.1, i.e., minimum read depth of 250 and minimum allele frequency of 10%) and “standard” (-v 100 -a 0.05) SNV calls. For mouse kidney WGS samples, standard calls were complemented with entries from the exhaustive output for variants with high confidence calls on at least one sample from the same strain.

For RNA-seq samples, allele frequencies were obtained from the exhaustive output for all variants in the corresponding strain WGS calls.

For WGBS and RRBS data, CpG methylation estimates were obtained from the bismark alignments using bismark_methylation_extractor on --paired-end or -‑single-end mode as appropriate. Alongside the alignments themselves and WGS variant calls from the same mouse strain or human subject, the per-read methylation calls that this tool generates were then used as input for blink, an in-house python tool for estimating allele-specific methylation and SNV frequencies, described more in detail below.

For Bis-PCR-seq data, only reads that mapped to the correct starting position and matched the consensus perfectly were used for further analysis. The R samtools interface Rsamtools was used to identify each read as having either an A or a C at position -104 and to determine the methylation status at position -133 of each read. Reads could therefore be assigned to either A_meth_, A_unmeth_, C_meth_ or C_unmeth_. The total number of reads in each group was summed and the proportion of A’s at -104 (A_meth_+A_unmeth_/A_meth_+A_unmeth_+C_meth_+C_unmeth_) and the methylation level at the -133 CpG site for A reads (A_meth_/A_meth_+A_unmeth_) was calculated for each sample.

**MEF ultra-long read Nanopore data processing and analysis.** Ultra-long read Nanopore libraries sequenced from MEFs were initially basecalled using albacore version 2.4.0 and aligned to the 18S and 28S regions from the BK000964.3 mouse rDNA reference using the minimap2 python interface mappy. From an original yield of 932,683 reads, only the 1760 mapping to any such regions were retained for further analysis. The raw signal fast5 files for these reads were later re-basecalled using guppy version 4.2.2 with the options specified in the dna_r9.4.1_450bps_modbases_dam‑dcm‑cpg_hac model configuration and template files. Generated fastq files were then aligned to the whole-genome plus rDNA mouse reference described above using minimap2 version 2.17 with parameters -c -ax map-ont. Reads aligning to the rDNA were then extracted and listed using samtools version 1.10.

Genetic variation and CpG methylation were estimated from the rDNA reads using megalodon on two different modes. In both cases, megalodon relied on a short-read variation information file adjusted using UpdateVCFSequenceDictionary from gatk version 4.1.6.0, used guppy 4.2.2 under the hood with res_dna_r941_min_modbases_5mC_CpG_v001 model configuration and template files from rerio, and produced per read modifications and variants (--outputs per_read_mods per_read_variants) in text format (‑‑write‑mods‑text and -‑write-variants-text), specifying CpG methylation as the target modification (-‑mod‑motif m CG 0). These output files vastly simplify the procedure necessary to obtain allele-specific methylation from Nanopore data compared to previous studies^46^. Even running on GPUs, it was necessary to increase the value of the ‑‑guppy-timeout option to 600 seconds to enable ultra-long reads to be successfully processed.

The first megalodon run was performed using version 2.2.6, with the strain-augmented variant file corresponding to Mouse 1 from the kidney C57BL/6J WGS samples and the same reference genome used for short-read alignments. Per-read variant calls were then analysed using an in-house R script in order to establish potential rDNA haplotypes. To this end, only calls with reported probability over 0.9 for either reference or alternative allele (but not both) were considered. For each pair of distinct positions, co-occurrence of allele combinations within reads were tallied, and such tallies then used as input for Fisher’s exact tests. The obtained p-values were then FDR-adjusted, and used as proxies for linkage strength between position pairs. Position -104 was chosen as the initial anchor point due to its well-established epiallelic nature. According to the tests, 1305 and 12736 proved the best linked positions to -104 overall and within coding regions, respectively. Although the linkage between -104, 1305, and 12736 appeared virtually perfect in reads with C at -104, exploration of allele frequencies across the remaining variant positions revealed that substantial variability remained in the “C” haplotype. Position 8063 was then found to maximally split the “C” haplotype into more consistent haplotypes, although some variability remained. Alleles at the four positions, obtained from reads with calls for all of them, formed four major combinations, plus a fifth one with a single read, which was discarded.

Individual reads were then assigned to their corresponding haplotype only if they had variant calls at all four positions. Since -104 and 1305 were perfectly linked, the allele at 1305 was not included in the names of the putative haplotypes, but all four positions were taken into account when assigning reads to haplotypes. 53 positions located upstream of -1000 and linked with either -104 or 12376 at a p-value < 10^-6^ were then considered as candidates for uniquely identifying the distinct haplotypes, as well as the two super-haplotypes defined by the allele at -104. Allele frequencies were calculated for each such position from reads assigned to each haplotype, keeping the 46 biallelic positions with minimum allele frequency smaller than 0.2 for all haplotypes. A position was then considered as specific to a haplotype if the frequency of one of its alleles exceeded 0.8 only in such haplotype (or super-haplotype).

The default megalodon execution above, however, can only analyze a single rDNA unit per read, regardless of how many the read actually spans, wasting much of the information that ultra-long reads contain. To address this issue, both the reference sequence and the input variation file were artificially expanded to comprise 20 back-to-back mouse rDNA units. Using these input files, megalodon version 2.2.9 was then executed with identical parameters as previously except for -‑allow‑supplementary-alignments and ‑‑reads‑per‑guppy-batch 10. The output per-read files of this multi-unit execution mode were then used to estimate haplotype-specific methylation and haplotype co-localization on the MEF.

Both per-read variant and modified bases estimates were first adjusted so that positions reflected their single-unit equivalent, and a unit number identifier was obtained according to where in the expanded reference they lay, distinguishing potentially-distinct units from each read. At the time of writing, however, megalodon is unable to distinguish between supplementary alignments from actually-distinct units or secondary alignments from the same unit to different positions of the expanded reference. To this end, multiple calls for the same reference position on the same read were considered as distinct only if their estimated probabilities (for reference and alternative allele, in the case of variants, and for canonical and modified base, in the case of methylation) differed. In addition, only calls with probability over 0.9 were accepted. This led to a total of 1750 rDNA units being represented across 1090 reads on the methylation calls, with a maximum of 11 units on a single read, and 1721 units across 1068 reads on the variant calls, with a maximum of 10 units on a single read. Units were then assigned to haplotypes based on their alleles at ‑104, 1305, 8063, and 12736, bringing the number of usable units down to 372 across 317 reads, with a maximum of 4 units on a single read. Haplotype frequencies remained largely unaltered compared to those obtained from megalodon’s default mode of operation (**Additional file 3: fig. S1**).

Haplotype co-localization frequency estimates were obtained on the 49 reads containing more than one haplotyped unit. The current implementation of megalodon does not preserve information about where in the read each call occurs, so the analysis focuses on haplotype co-appearance across entire reads and not necessarily physical contiguity of the haplotyped units. In particular, for any given haplotype H1, the average proportion of units of each haplotype H2 (which might be the same as H1) was calculated on reads containing at least one H1 unit.

Haplotype methylation estimates, at both CpG site and coding unit level, were also obtained from the multi-unit megalodon calls. For the former, a mean methylation level was computed across all units assigned to each haplotype for CpG sites if more than 10 units of the haplotype had methylation calls for such a site. Promoter and transcribed region estimates were calculated on sites located between -1000 upstream of the TSS and the end of the 3’ETS, whereas IGS estimates were obtained on sites downstream of the 3’ETS and up to the end of the looped rDNA unit reference (-3008 upstream of the TSS). Methylation at the coding unit level was estimated as the mean methylation level across all CpG sites between -1000 upstream of the TSS and the end of the 3’ETS for each haplotyped unit separately.

**blink: Allele-specific methylation and frequency from bisulfite-converted data.** An in-house python tool called blink was developed to jointly estimate allele-specific methylation levels and allele frequencies from bisulfite-converted short-read sequencing data. In particular, blink takes as input the bismark alignments of a sample, the output of bismark’s bismark_methylation_extractor tool (hereinafter, “bme files”), and SNV calls from WGS data in vcf format. Alignments and variant calls are processed under the hood using pysam, the python interface to samtools. Original Top (“OT”) and Original Bottom (“OB”) bme files are processed separately, adjusting the position of the methylation calls in the latter to coincide with those in the former. Base calls at variant positions are extracted from the alignments, tallying them separately according to in which bme file their original reads appear. Since bisulfite conversion conflates different bases depending on their direction, Cs and Ts are tallied jointly on OT reads, and As and Gs on OB reads.

In blink’s default mode of operation, intended for WGBS data, only positions covered by a given minimum of reads from each bme file are kept, and allele-specific counts are obtained solely from the direction in which they can be distinguished, ignoring any that does not exceed a given minimum frequency threshold. Unless a “multiallelic” mode is explicitly requested, only the two most frequent alleles are considered believable and kept for further analysis. In RRBS data, however, alignments tend to be highly directional, with variant positions often solely covered from OT or OB reads. Highly imbalanced alignments hamper the ability to distinguish between conflated bases, so blink offers an “informed” mode of operation in which the minimum required coverage is considered jointly for both bme files, and alleles are inferred based on those appearing on the input SNV file. C/T variants covered exclusively from OT reads, however, as well as A/G variants covered exclusively from OB reads, cannot be disentangled and thus are removed from further analysis. Moreover, this “informed” mode might introduce biases in the analysis and it is thus only recommended for RRBS data.

For those variant positions in which alleles can be inferred, blink then imputes the most likely allele to those reads with conflated bases, correcting for the bisulfite conversion, and estimates allele frequencies based on such imputations. These frequencies are provided alongside those from the input SNV file for comparison. Finally, blink joins corrected variant calls with CpG methylation calls on a per-read basis, producing a data structure with one entry for each successfully-called SNV and CpG site pair appearing on each input read.

The source code for blink is available online at https://github.com/franrodalg/blink.

In this study, blink was run for all bisulfite-converted datasets with minimum coverage of 10 reads and minimum allele frequency of 0.01, on “default” operation mode for WGBS data and “informed” for RRBS data. For the particular case of the mouse kidney WGBS data, blink was run for each sample using their corresponding strain-complemented SNV lofreq calls described above. For all RRBS public datasets, blink was provided with the strain-complemented SNV lofreq calls from C57BL/6J Mouse 1 kidney. Finally, for the human LCL WGBS data, blink was run on “multi-allelic” mode (i.e., allowing for more than two alleles per position) using the variant calls from the corresponding publicly-available WGS sample.

**Haplotype-specific analysis of C57BL/6J kidney and muscle data.** In order to assign reads to haplotypes, the blink output for the C57BL/6J kidney WGBS data was matched with the haplotype-specific alleles identified from the MEF ultra-long Nanopore data according to SNV position and called allele. For each sample, only SNV-CpG site pairs with at least 10 reads per haplotype, and 50 overall, were considered for computing haplotype-specific methylation levels as the proportion of positive methylation calls over all reads assigned to the haplotype covering each CpG site.

Predicted ‘A’-associated methylation levels in **fig. S3** from **Additional file 3** were calculated for each C57BL/6J kidney sample using the following formula:

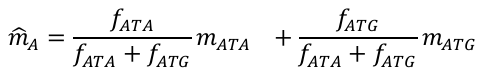

where $f_{ATA}$ and $f_{ATG}$ are the average allele frequencies from WGS at haplotype-associated SNVs, and $m_{ATA}$ and $m_{ATG}$ are the average methylation levels from WGBS at haplotype-associated CpG sites.

For non-bisulfite-converted data (WGS, mRNA-seq, rRNA-seq, and CUT&Tag), haplotype-specific frequencies were obtained by matching the entries from lofreq variant calling files with the MEF ultra-long Nanopore haplotype-specific alleles according to their position. The frequency reported for each position was then adjusted to its complementary value if the haplotype-specific allele corresponded to the reference, and left untouched otherwise.

**Processing and analysis of C57BL/6J publicly-available datasets.** For all analyzed datasets, reads were assigned to haplotypes following the same procedure described above for kidney and muscle data, and haplotype-specific methylation calculated using the same thresholds.

In the particular case of the ChIP-Seq data from Diesch *et al*.^17^, overall haplotype frequencies for each replicate and condition were obtained by summing the total depth corresponding to the haplotype and dividing it by the total depth at the associated positions. Relative frequencies were then calculated dividing the haplotype-specific frequencies from UBTF samples by their equivalent from the input samples.

Per-sample haplotype-specific methylation levels for environmental exposure^6,16^, Dnmt1 KO^17^, and ageing^19^ were calculated as the proportion of positive methylation calls over all haplotype-associated sites. Reported p-values for the difference of means were obtained using Wilcoxon rank sum tests.

rDNA presence was checked in all mRNA-seq samples from Dahlet *et al*.^17^ using samtools view -c, obtaining between 3.2 and 14.3 million reads mapping to the rDNA reference. Relative ATA- and ATG-associated frequencies were calculated similarly to the haplotype-specific frequencies from Diesch *et al*. above, but only considering ITS2 variant positions to avoid differential rRNA depletion between coding subunits and transcribed spacers affecting. Minimum coverage at the considered positions was 388 reads, as reported by samtools depth, with most samples being in the thousands. The reported p-value for the difference of means was also obtained using a Wilcoxon rank sum test.

Correlation between age and methylation from Petkovich *et al*.^19^ was calculated on samples labelled “C57BL/6” from ages 3 months and above. For each age group, CpG sites covered by fewer than 3 samples were discarded, and only those sites that appeared in at least 5 age groups were further analysed. Methylation levels from each remaining site were correlated with age using Pearson’s method. Due to the relatively low RRBS coverage at ATG-associated positions, estimates for such haplotype were obtained on fewer than 3 CpG sites per sample on average. All other haplotypes retained at least 15 CpG sites on average.

The “expected” correlation value for the ATA haplotype was obtained assuming that methylated units lose methylation at the same pace *R* as unmethylated units gain it. If that was the case, the observed correlation coefficient for the non-ATA haplotypes would be:

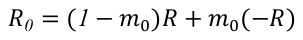

with $m_{0}$being the observed overall methylation of non-ATA haplotypes at 3 months. Isolating $R$ from the equation above yields:

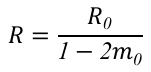

If ATA units lost (or gained) methylation at the same pace, their associated correlation coefficient would be:

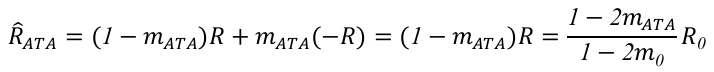

Substituting with the precise values for this dataset ($m_{ATA}$=0.752, $m_{0}$=0.235, and $R_{0}$=0.34) leads to an “expected” correlation coefficient for ATA of approximately $\hat{R}_{ATA}$=-0.32.

The degree of disorder in samples from Petkovich *et al*.^19^ was estimated using two different methods: per-read entropy and epipolymorphism. In the former, only reads containing at least 10 CpG sites were considered. Shannon entropy was then calculated for each read, using the proportion of methylated sites as probability value. These entropies were averaged per haplotype for each sample and correlated with age using Pearson’s method. Epipolymorphism, a disorder metric first introduced by Landan *et al*.^27^, was implemented using the method Scherer *et al*. detail^29^. In particular, the four CpG sites associated with the SNV at 6832 appearing in the highest number of reads and in all samples were selected. Reads missing methylation calls for any of the four sites were then removed, and the remainder were used to estimate the proportion of each combination of methylation states appearing per haplotype and sample. The corresponding epipolymorphism values were then calculated as 1 minus the sum of all proportions squared. These values were then correlated with age in the same way as the per-read entropy above.

**Total rDNA copy number analysis.** Total rDNA copy number estimates were obtained using orthologous methods: droplet digital PCR (ddPCR) and short-read sequencing (WGS and WGBS).

Estimates from ddPCR were obtained from the Quantasoft software in default mode. These estimates are derived from the fraction of droplets with positive fluorescence detection of the target 18S amplicon using a Poisson statistic.

On the other hand, estimates from short-read sequencing data (WGS and WGBS) were obtained following the procedure Gibbons *et al*^44^ suggest. In particular, reads were aligned to the exome references described above, obtaining the average read depth per sample from the output of samtools depth. These values were used as proxies for single-copy gene coverage in order to normalise the average read depth at the 18S subunit from whole-genome plus rDNA alignments. rDNA copy number estimates were then calculated as twice the quotient between 18S and exome average read depth.

**Analysis of mouse strains kidney data.** Allele-specific methylation levels were obtained at each CpG site from blink output files using the same coverage threshold as indicated above for haplotype-specific analyses. For each SNV, a p-value for the difference in mean CpG methylation between alleles was obtained using a Wilcoxon rank sum test. These p-values were then FDR adjusted at sample level. SNVs were considered as candidate epivariants, and displayed in **Fig. 4A**, if at least 3 samples in a strain had allele-specific methylation calls there, and the methylation differences were significant at FDR < 0.01 for over half of those samples.

Allele frequencies at the candidate epivariant positions were extracted from the lofreq output files for both WGS and rRNA-seq data. Methylation-adjusted WGS alternative allele frequencies were then computed as follows:

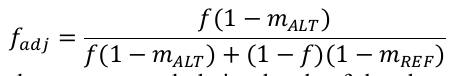

where $m_{ALT}$and $m_{REF}$ are the average methylation levels of the alternative and reference allele, respectively, and *f* represents the alternative allele frequency.

rDNA promoter and 18S methylation levels used to analyse association between rDNA copy number and methylation were obtained from the bismark_methylation_extractor output, using positions between -1000 and the TSS as proxies for the rDNA promoter. In the case of exon methylation, CpG methylation values were extracted from the bismark reports generated from exome reference alignments. All correlation estimates were obtained using Pearson’s method.

**Analysis of human LCL data.** Allele-specific methylation levels were estimated from the blink output for each of the 48 human cell lines using the same coverage thresholds described above. Wilcoxon rank sum tests for differences in mean methylation were computed for SNVs covered by at least 24 samples. FDR-adjusted p-values for positions -413 and 7980 from the KY962518.1 rDNA reference were lower than 0.01, and were thus selected for further study. Variation at -413 was observed in 38/48 samples, whereas 7980 showed variation in all considered samples. The average methylation level around each position was obtained as the weighted mean between the methylation levels of their corresponding alleles. rDNA copy number values were estimated from publicly-available WGS data from the 1000 genomes project^32^ as described above for the mouse. These estimates highly correlate (Pearson’s p-value < 2.2x10^-16^) with those Hall *et al*.^31^ provide for the same individuals (**Additional file 3:** **fig. S19**). Since -413 is located in the promoter, its observed RNA expression did not reach minimum depth threshold and was thus discarded from further study. Allele frequencies at 7980 were estimated from WGS and rRNA-seq data using lofreq and adjusted for methylation following the same procedure as above.

**Human ultra-long read Nanopore data processing and analysis.** Publicly available MinION ultra-long read Nanopore data for human sample HG02723 was obtained from Circulomics (<https://www.circulomics.com/datasets>). This sample corresponds with a Mandinka female offspring individual (<https://www.internationalgenome.org/data-portal/sample/HG02723>) from the 1000 genomes project. This sample was not included in the LCL analysis above since offspring cell lines are not available to acquire from the Coriel lab. Raw fast5 files were basecalled using guppy version 4.2.2 as above, yielding a total of 459,549 reads. These reads were then aligned with minimap2 to the human whole-genome plus rDNA reference sequence described above. A total of 1,233 reads aligned to the rDNA and were considered for further analysis with megalodon version 2.2.9, using a reference sequence artificially expanded to include 20 back-to-back rDNA units and the same parameters described for the MEFs. Short-read SNVs were obtained by first reconstructing with samtools fastq the raw sequencing reads from the CRAM file available from the 1000 genomes project at <ftp://ftp.sra.ebi.ac.uk/vol1/run/ERR398/ERR3989060/HG02723.final.cram>, and then trimming, aligning and variant calling as previously described. These SNVs were then used as input in megalodon to obtain per-read variant and CpG methylation calls.

Genetic linkage was computed using Fisher’s exact test between every variant pair in the rDNA “coding unit”. The most closely linked positions were then selected as candidates for haplotype analysis, first using 7980 as anchor point (note: this sample did not display variation at -413), and later extending to any set of variant positions. Following the same procedure as described for the MEF data did not yield any combination of variants with clear haplotypic structure. For instance, positions 6521, -348, and 12986 were the top linked variants with 7980 overall (p-value = 4.71x10^‑24^), in the promoter (p-value = 3.72x10^-14^), and in the 28S (p-value = 5.70x10^-12^), respectively. 112 rDNA units across 95 reads had variant calls in all four positions, yielding nine observed combinations of alleles (**Additional file 3:** **fig. S20**, **top panel**). Of these, only four appeared in more than 5 rDNA units: CCAC, CCGC, CGGC, and CGGT. Allele frequencies for the other coding unit variant positions, however, retained a substantial degree of variability after assigning units to the putative haplotypes, even when only considering the four listed previously (**Additional file 3:** **fig. S20**, **bottom panel**). Other combinations of SNVs, regardless of whether 7980 was used as anchor or not, yielded similarly variable allelic structures.

Methylation values for **Fig. 5D** were calculated as the mean of the CpG methylation calls between position -1000 and the end of the 3’ETS for each individual rDNA unit with variant call at 7980 and spanned at least from the promoter (i.e., had CpG sites called upstream of the TSS). At the time of writing, megalodon does not retain spatial information in its supplementary alignments mode when the alignments are split. This means units adjacent in a molecule might appear as disjointly aligned. Variant calls from a particular unit might also miss due to quality thresholds not being met. **Fig. 5D** displays gaps between these potentially-disjoint calls.
