## Additional File 3 - Supplementary Figures for "Genetic variation at mouse and human ribosomal DNA influences associated epigenetic states"

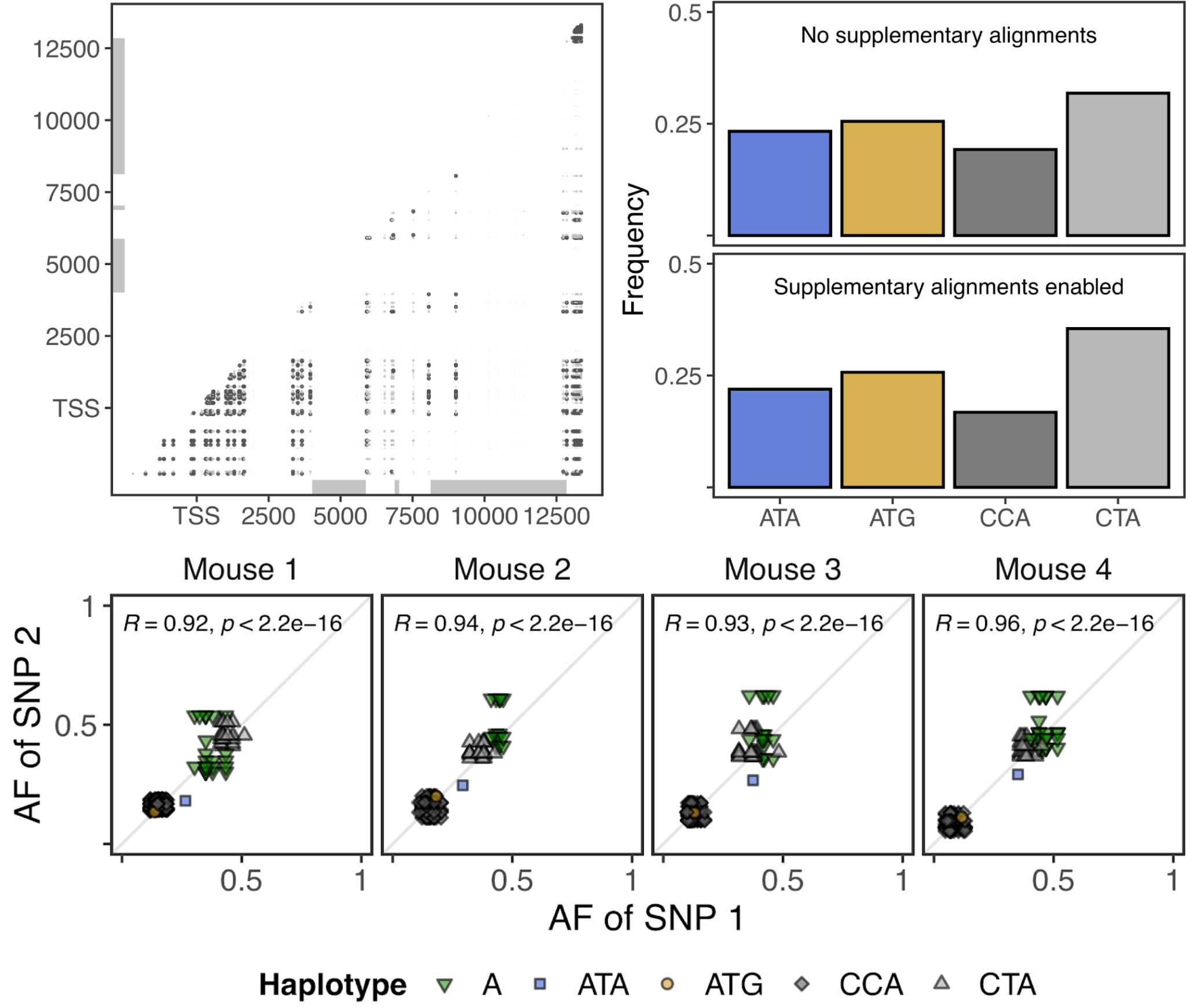

### **Figure S1. Identification and validation of C57BL/6J rDNA haplotypes.** Significantly-linked position pairs (Fisher’s exact test, FDR < 0.01) on the MEF ultra-long read Nanopore data (**Top left**). Observed frequencies for each rDNA haplotype before and after enabling supplementary alignments on megalodon (**Top right**). Correlation between allele frequencies in kidney C57BL/6J WGS data between pairs of haplotype-specific SNV positions (**Bottom**). Points corresponding to the same haplotype tend to cluster together.

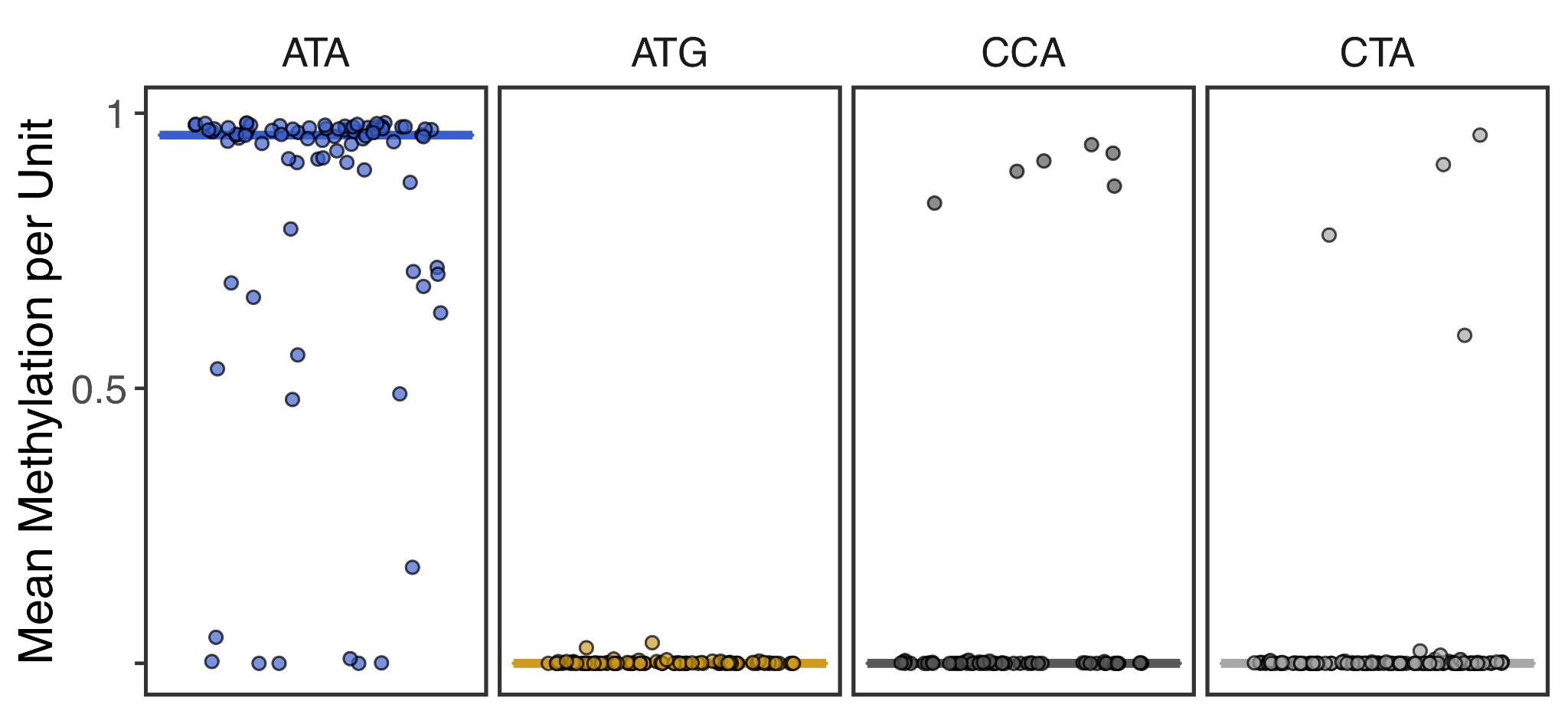

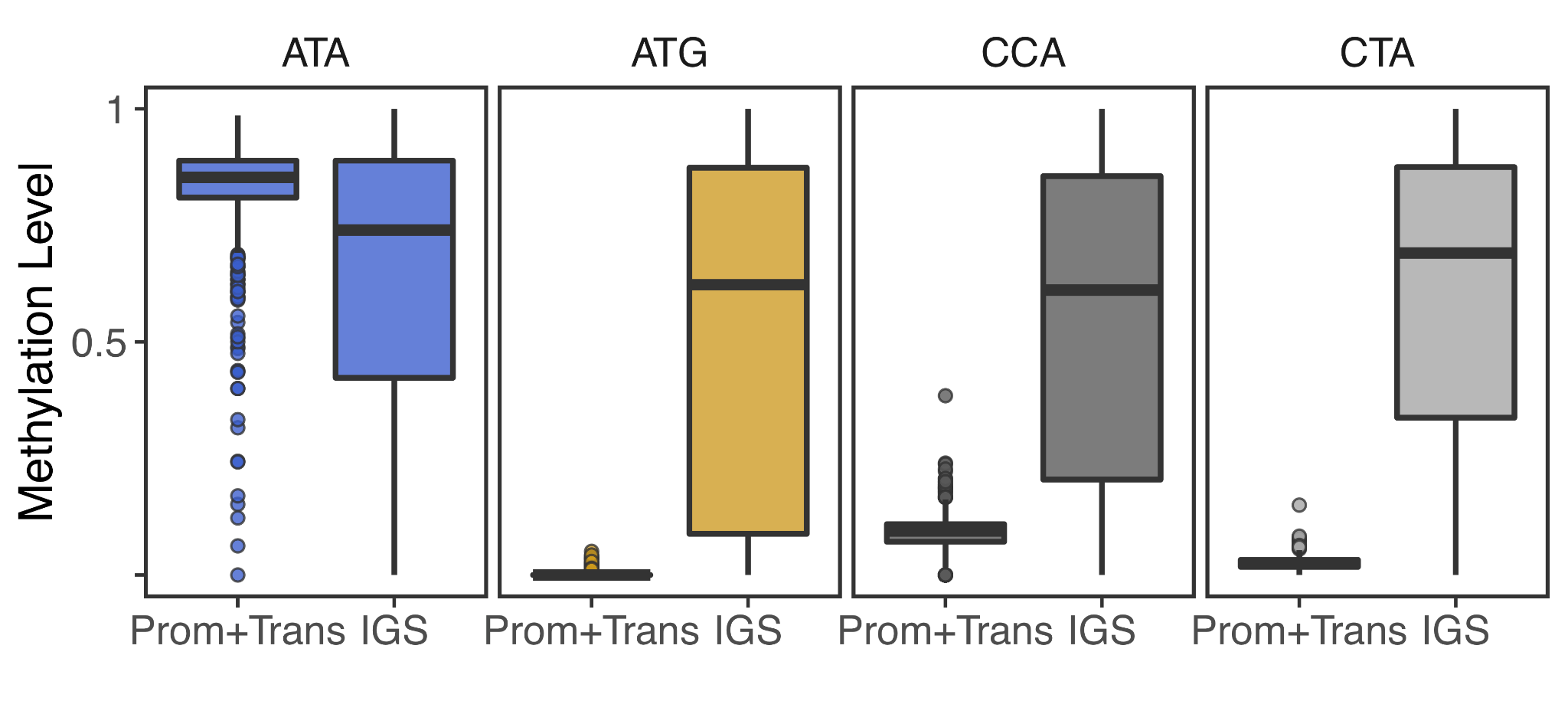

**Figure S2. Haplotype-specific methylation in C57BL/6J rDNA from Nanopore data.** Average methylation levels per unit (**Top**), and comparison between methylation levels at promoter and transcribed rDNA regions (the “coding unit”), and the IGS (**Bottom**), for the C57BL/6J rDNA haplotypes in MEF ultra-long read Nanopore data.

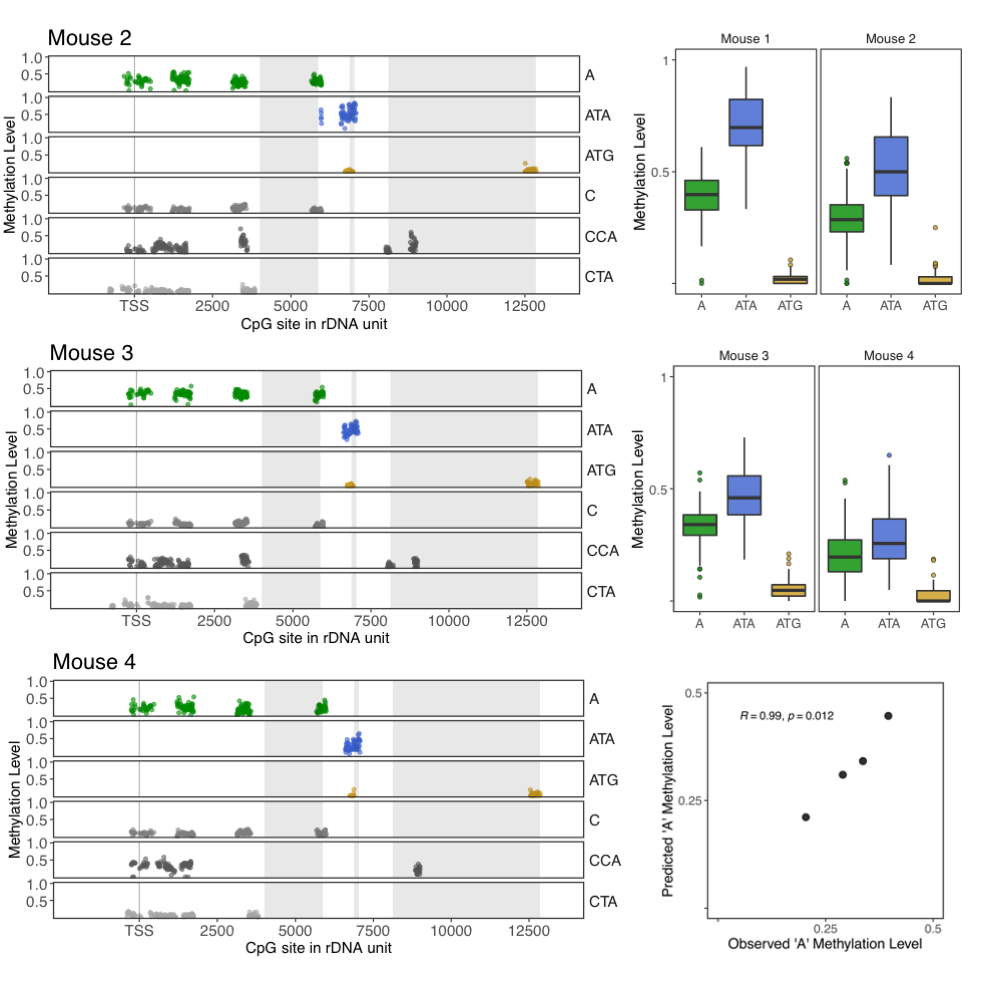

### **Figure S3. Methylation levels per rDNA haplotype in kidney C57BL/6J WGBS.** Positional data for the mice not shown in **Fig. 2B** (**Left**), boxplot summaries for the ‘A’-related haplotypes in the four kidney C57BL/6J WGBS samples (**Top- and middle-right**), and correlation between methylation observed at ‘A’-associated CpG sites and ‘A’ methylation predicted as weighted average of ATA and ATG methylation levels (**Bottom-right**).

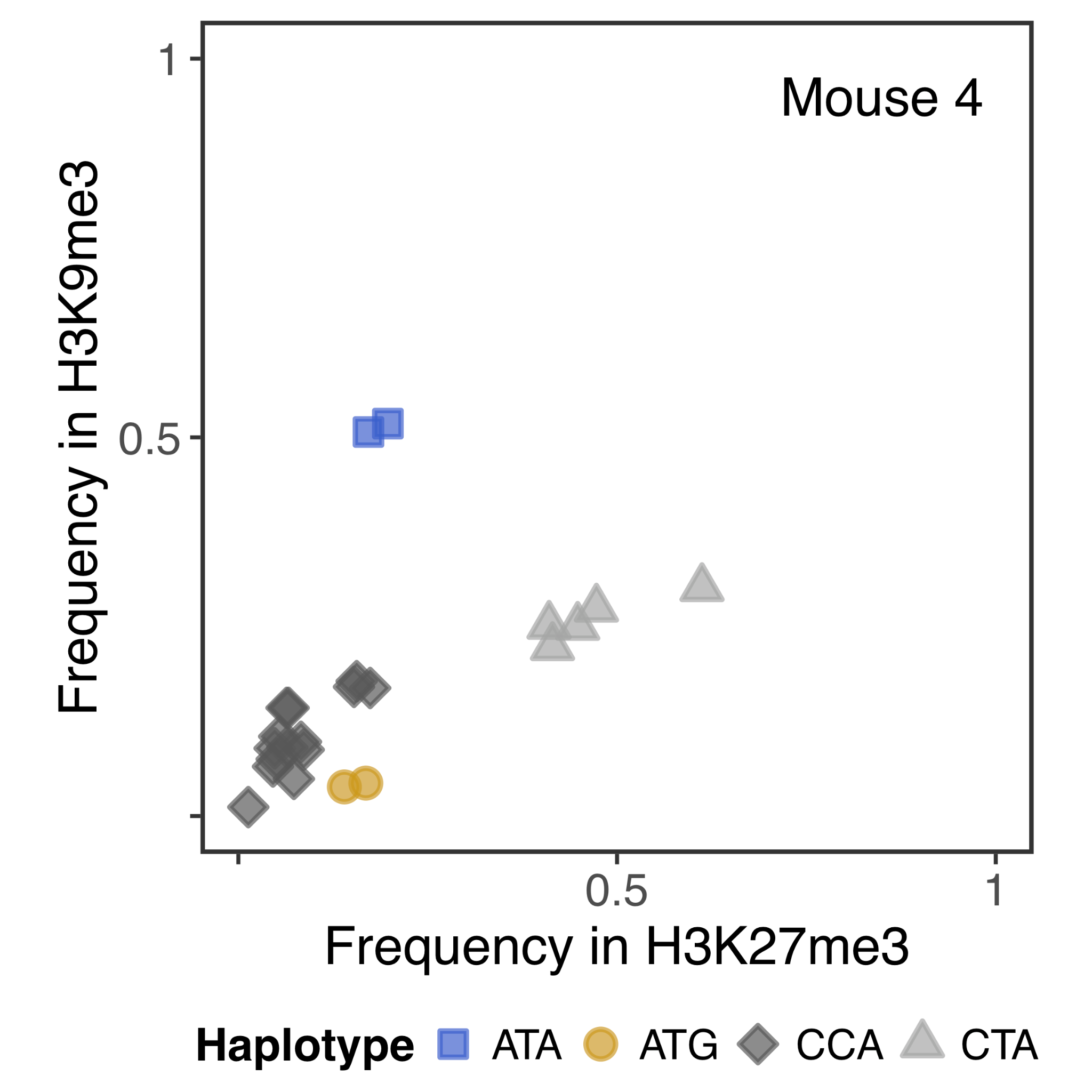

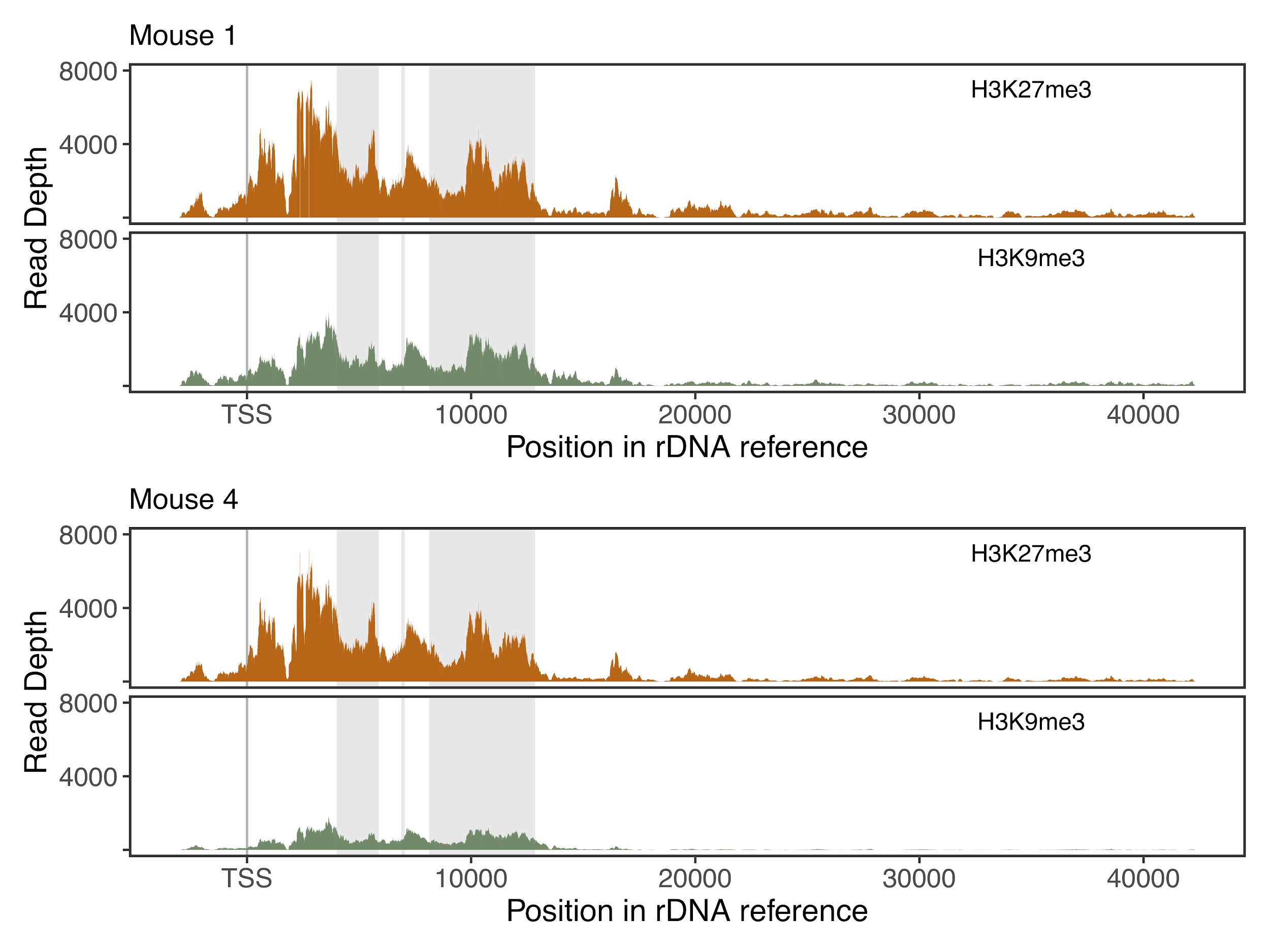

### **Figure S4. C57BL/6J kidney CUT&Tag data.** Comparison of haplotype-specific allele frequencies at variant positions between H3K27me3 and H3K9me3 reads (**Top**), and read coverage distribution across the rDNA unit (**Bottom**).

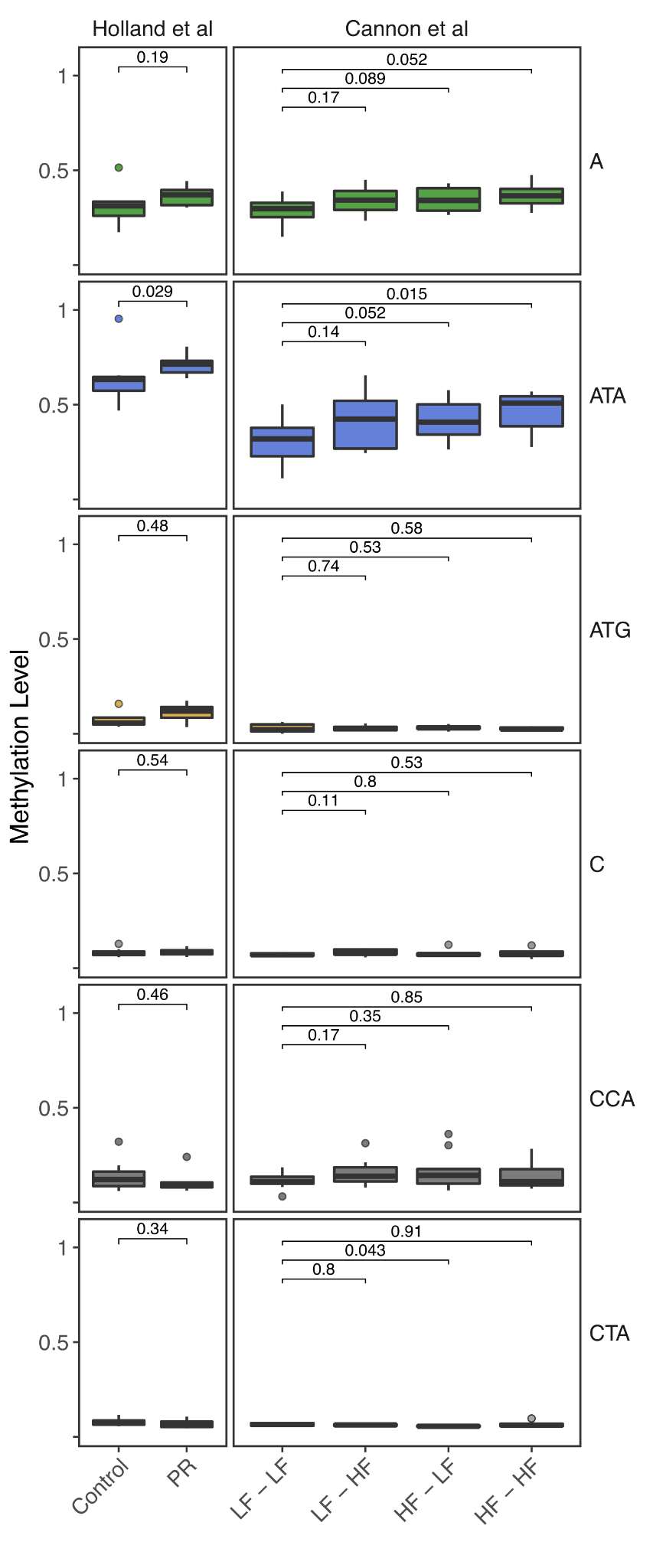

### **Figure S5. Haplotype-specific methylation in nutritional stress C57BL/6J datasets.** Extended version of **Fig. 3A** with non-ATA methylation levels separated by haplotype, and the LF-HF condition from Cannon *et al*.^16^ included. (Note: although the comparison between the LF-LF and HF-LF conditions for CTA haplotypes is associated with P = 0.043, the actual methylation difference is 0.0081%, i.e. a biologically meaningless difference. This is a result of the non-parametric tests used, which do not take into account the actual methylation difference.)

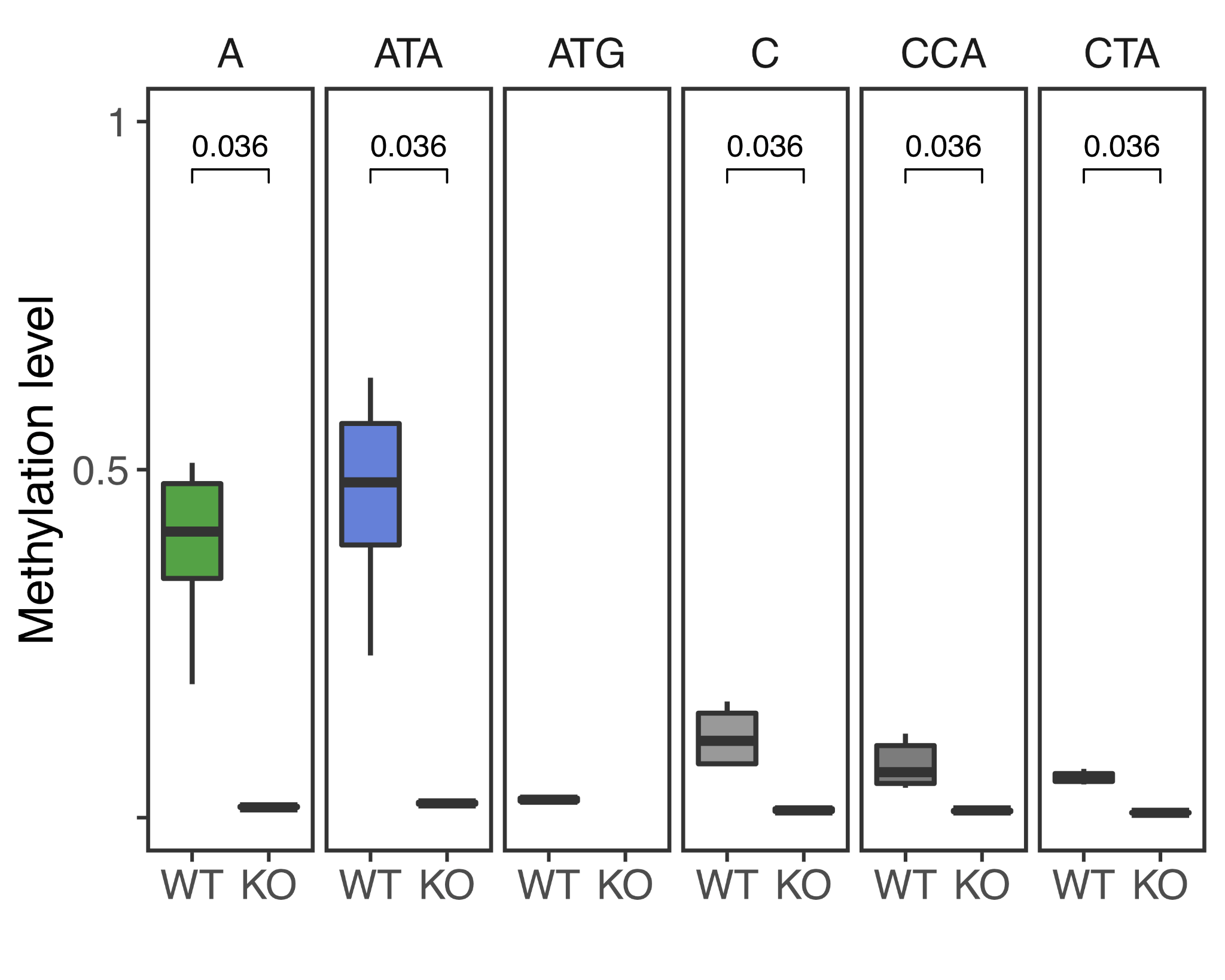

### **Figure S6. Haplotype-specific methylation levels on wild-type (WT) and Dnmt1 KO (KO) mouse embryos.** Methylation levels from RRBS data significantly decrease for all detectable rDNA haplotypes in Dnmt1 KO 8.5dpc embryos from Dahlet *et al*.^17^

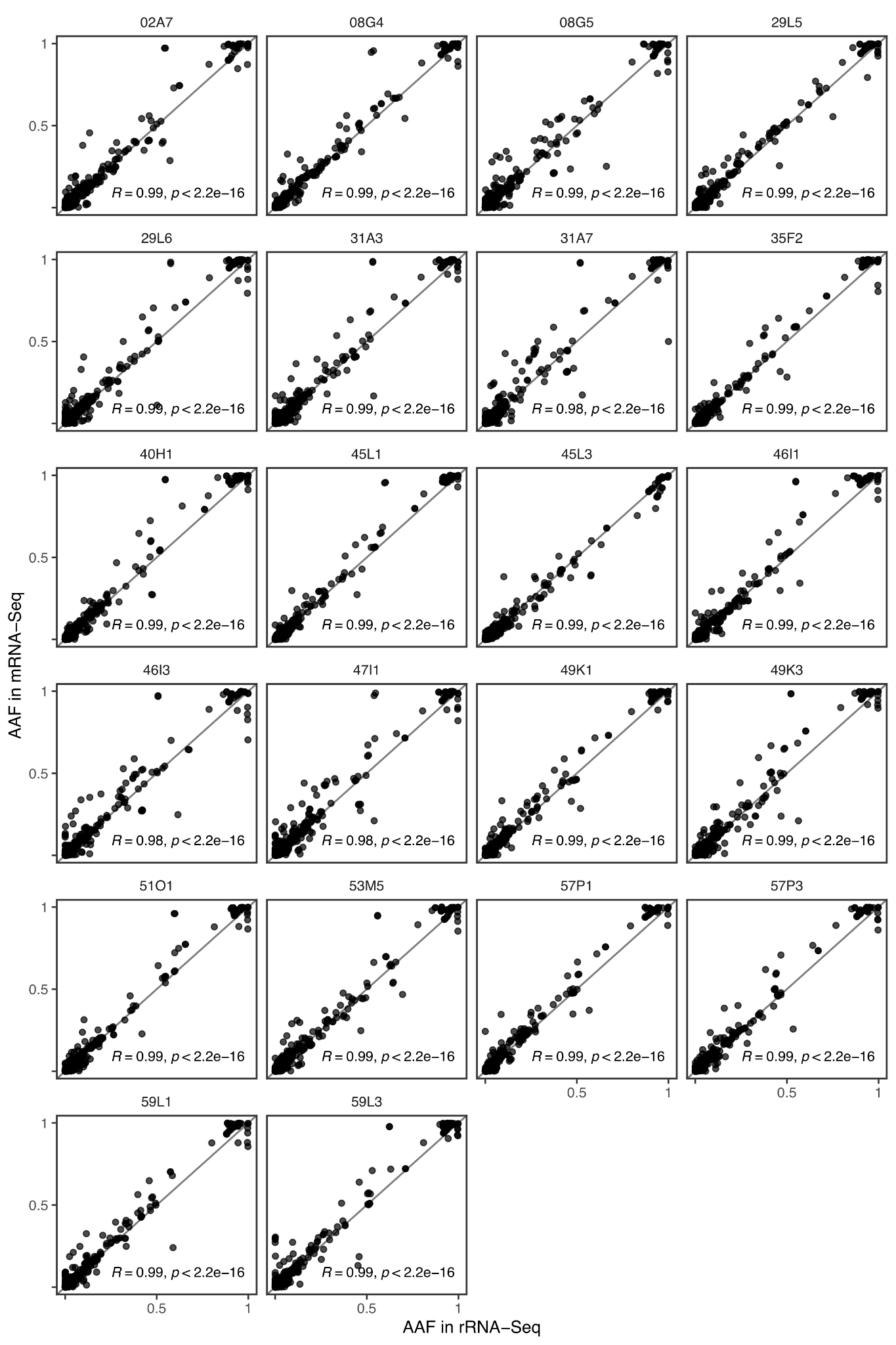

### **Figure S7. Association between mRNA-seq and rRNA-seq allele frequencies on C57BL/6J muscle.** Pearson correlation coefficients and p-values calculated on rDNA variants suggest alternative allele frequencies (AAF) closely match regardless of whether rRNA is depleted.

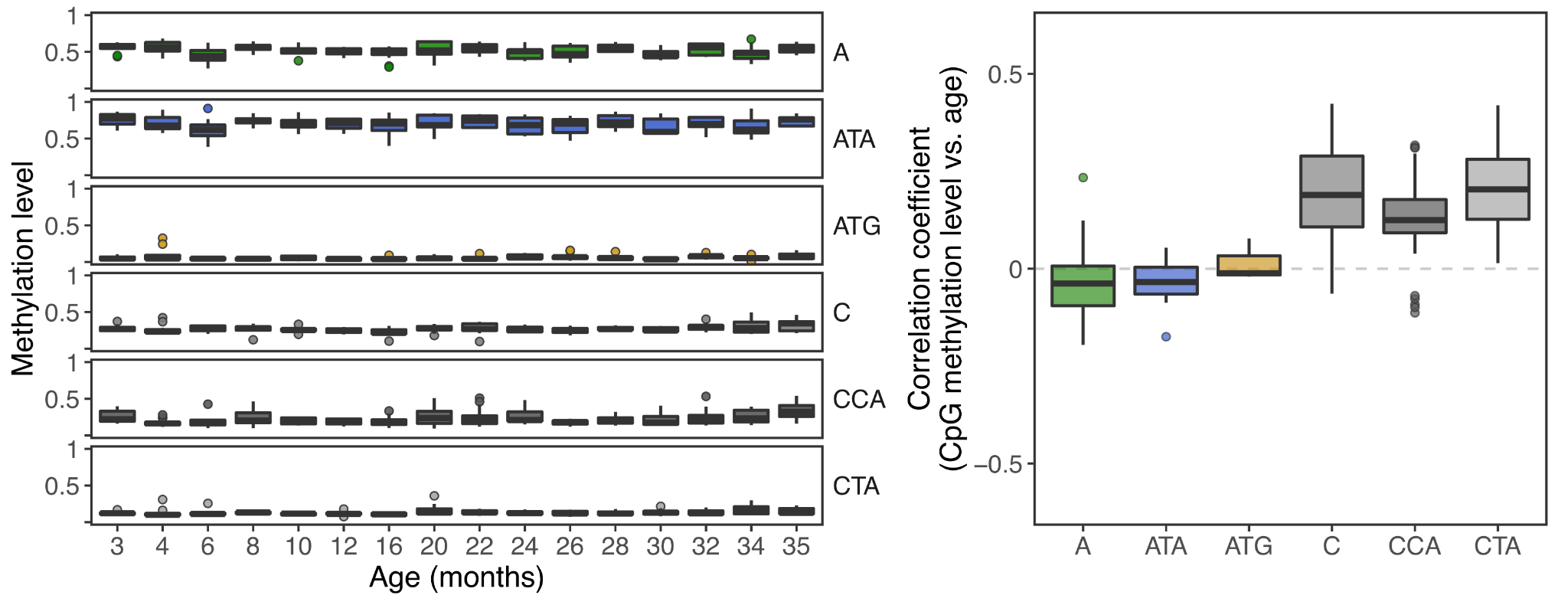

### **Figure S8. Haplotype-specific methylation across age groups.** Methylation levels (**Left**) and Pearson correlation coefficients (**Right**) obtained for the rDNA haplotypes in RRBS data from Petkovich *et al*.^19^ Estimates for ATG are obtained from fewer than 3 CpG sites per sample, on average, whereas estimates for the other haplotypes rely on an average of 15 or more CpG sites per sample.

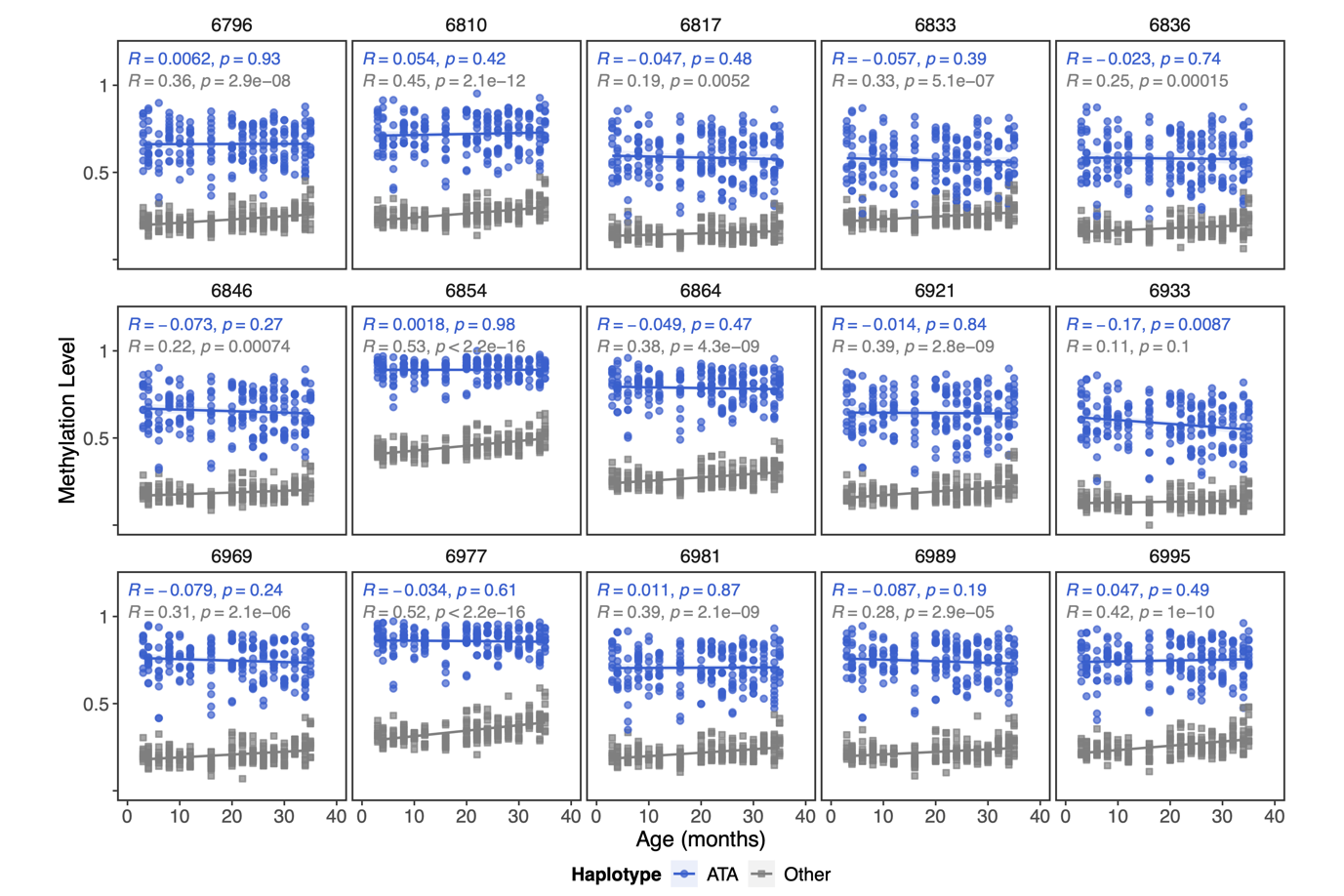

**Figure S9. Changes in methylation levels at ATA-associated CpG sites with age.** Only non-ATA haplotypes display statistically-significant accumulation of methylation in 14/15 CpG sites neighbouring the ATA-associated SNP at position 6832 in RRBS data from Petkovich *et al*.^19^

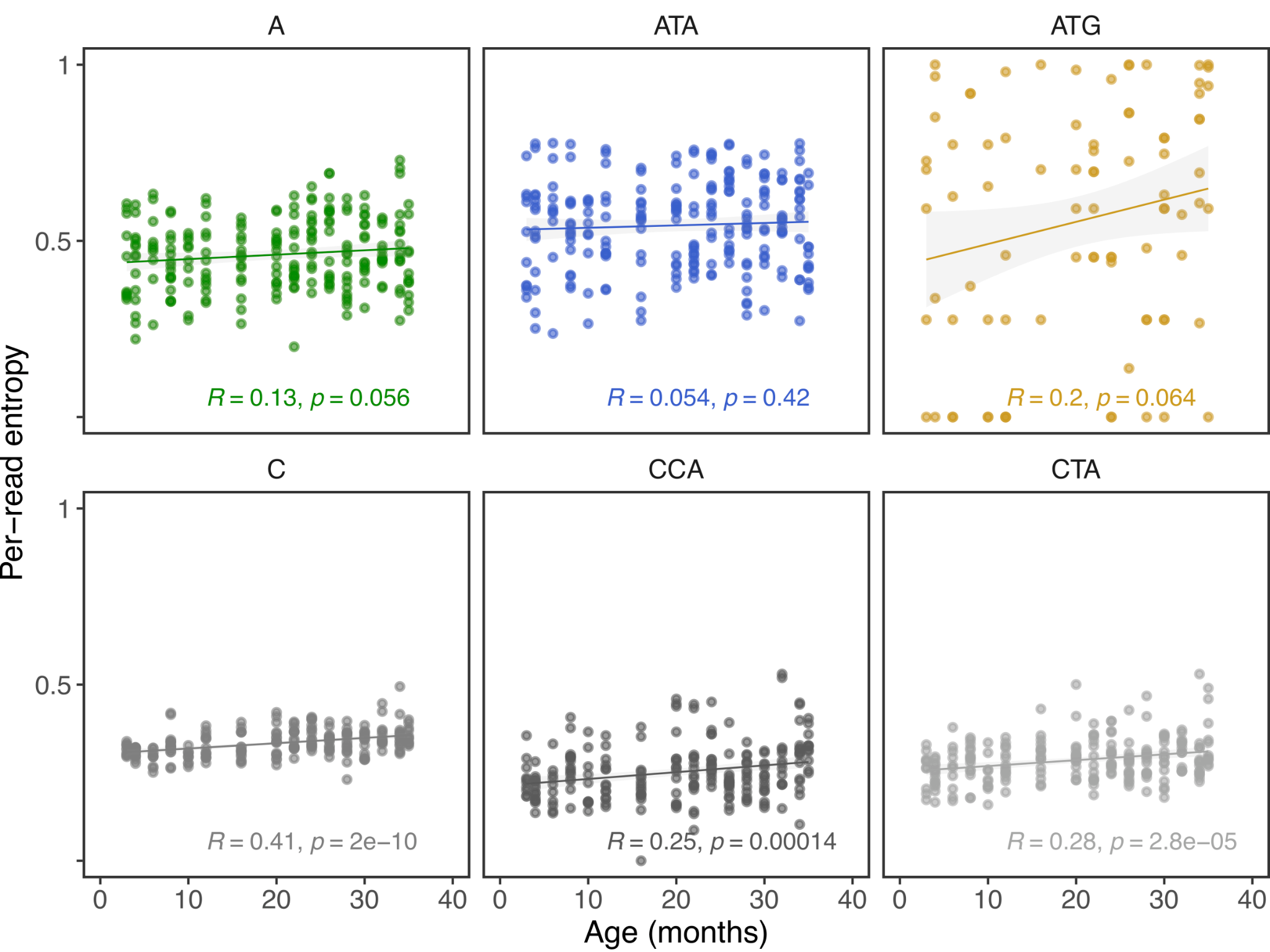

### **Figure S10. Haplotype-specific correlation between per-read entropy and age.** Extended version of **Fig. 3D**, with per-read entropy computed for all rDNA haplotypes in RRBS data from Petkovich *et al*.^19^ Estimates for ATG are obtained from fewer than 3 CpG sites per sample, on average, whereas estimates for the other haplotypes rely on an average of 15 or more CpG sites per sample.

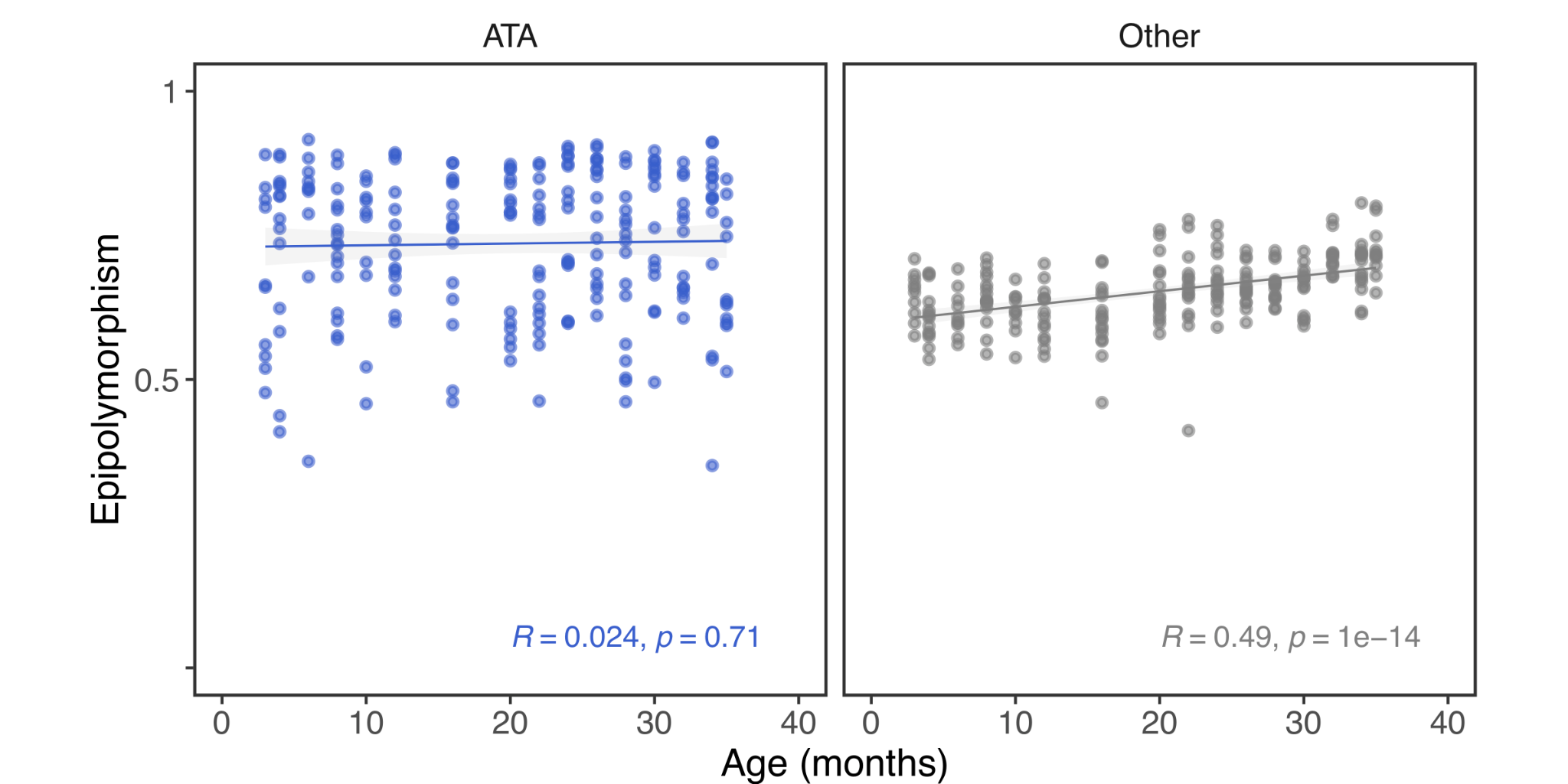

### **Figure S11. Association between epipolymorphism and age.** Similar to the per-read entropy in **Fig. 3D**, only non-ATA haplotypes display increasingly-disordered DNA methylation profiles with age in RRBS data from Petkovich *et al*.^19^

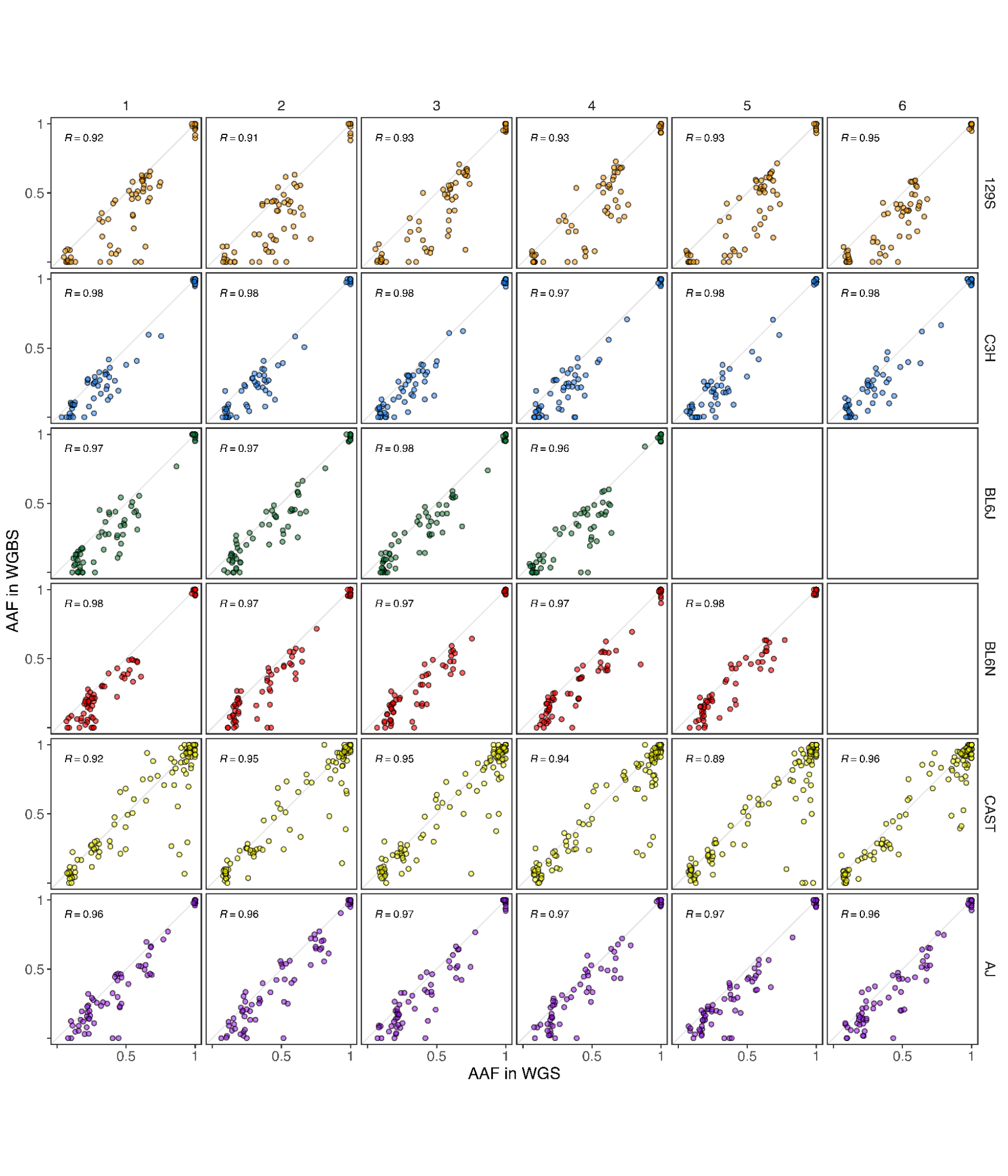

### **Figure S12. Association between estimated WGS and WGBS alternative allele frequencies on kidney mouse samples.** Pearson correlation coefficients between alternative allele frequencies (AAF) obtained from lofreq for WGS and the blink in-house script for WGBS, calculated on mouse kidney data from six different strains: 129S1/SvImJ (‘129S’), C3H/HeJ (‘C3H’), C57BL/6J (‘BL6J’), C57BL/6N (‘BL6N’), CAST/EiJ (‘CAST’) and A/J (‘AJ’).

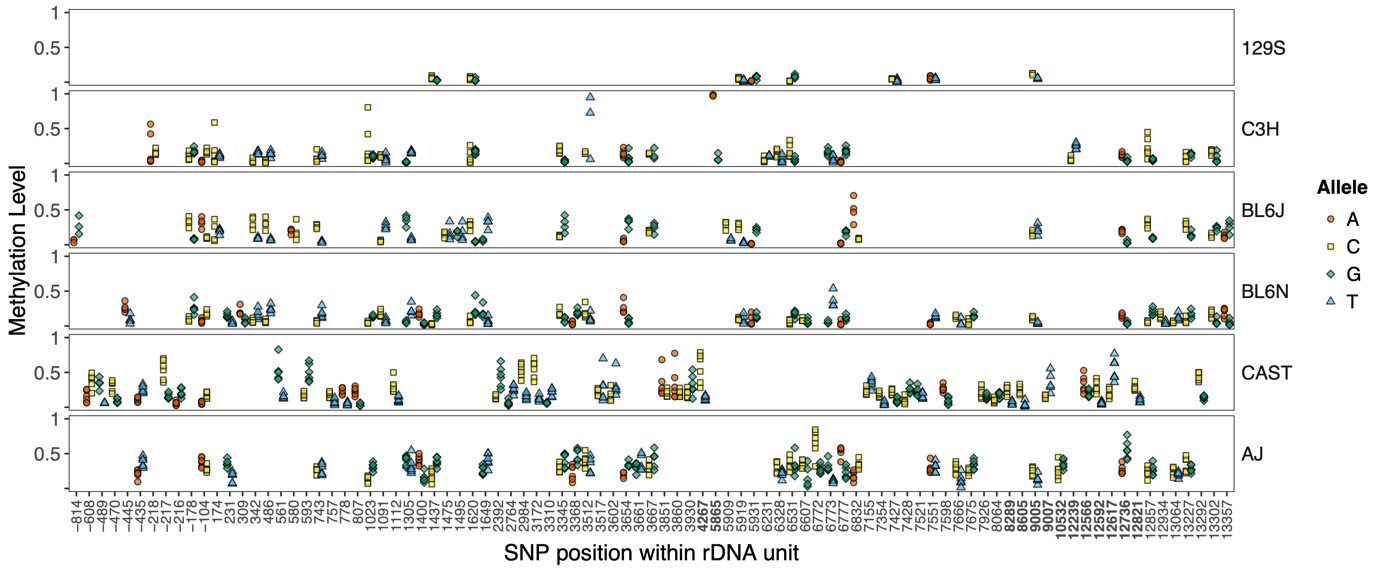

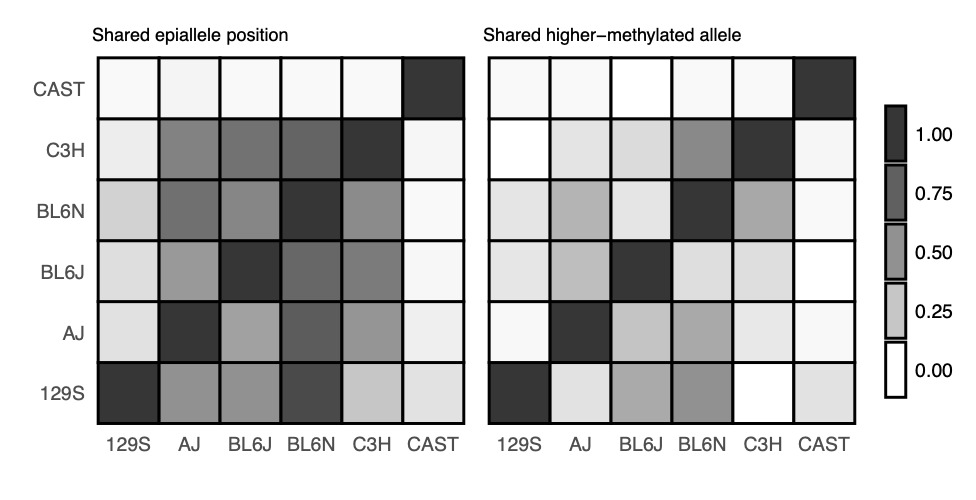

### **Figure S13. Comparison between epivariants across strains on mouse kidney samples.** (**Top**) Identification of epivariants in kidney of 5 inbred strains, C57BL/6J (‘BL6J’): 129S1/SvImJ (‘129S’), C3H/HeJ (‘C3H’), C57BL/6N (‘BL6N’), CAST/EiJ (‘CAST’) and A/J (‘AJ’). For each strain, displayed positions show statistically significant allele-specific methylation differences (FDR-adjusted p-value < 0.01) for over half of the mice. (**Bottom**) Concordance of epivariants across strains. Each cell represents the proportion of epivariants of the strain in the x-axis that also appear as epivariants in the y-axis strain, with the **left** panel only considering position concordance and the **right** panel also requiring the higher-methylated allele to match.

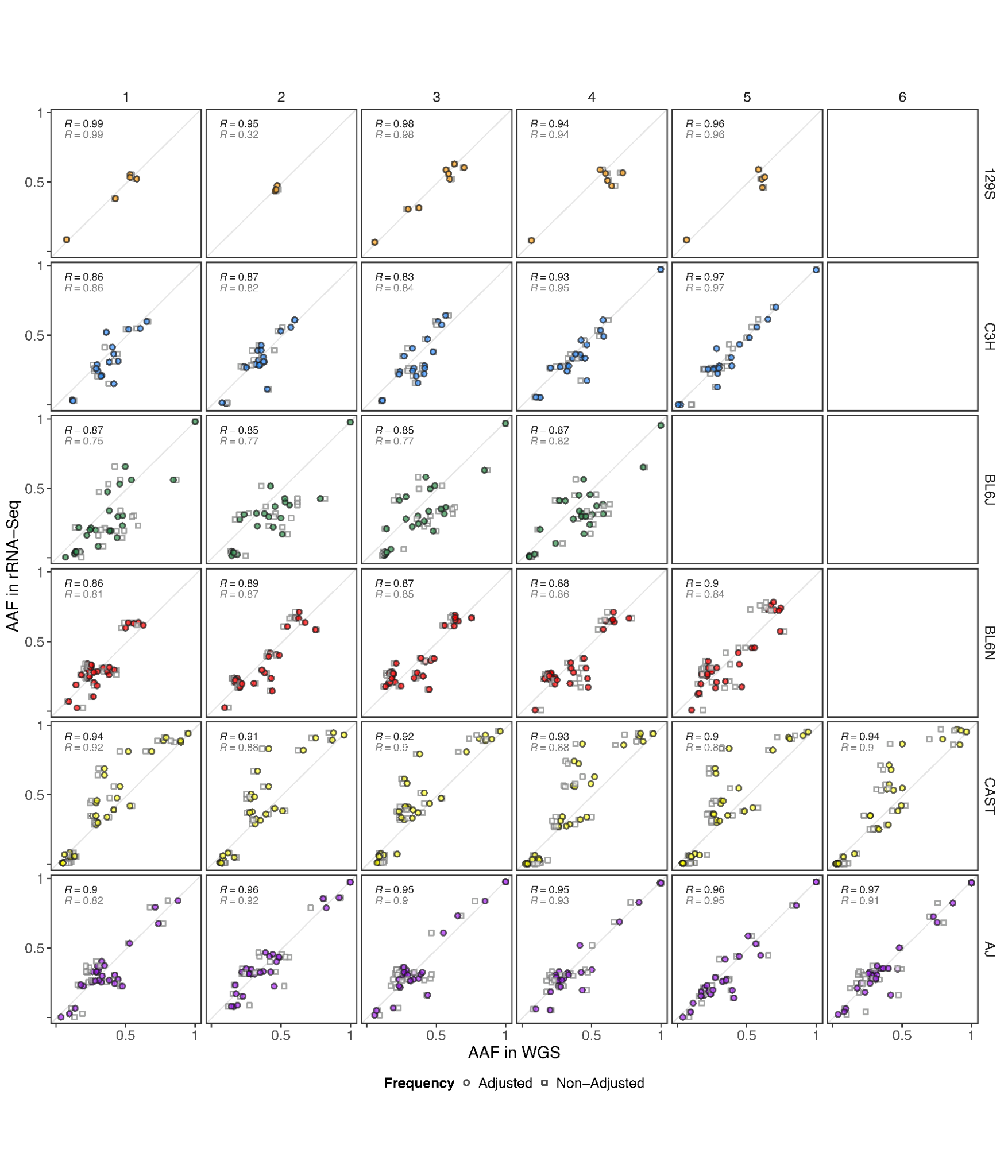

### **Figure S14. Association between WGS and rRNA-seq alternative allele frequencies on kidney mouse samples.** Pearson correlation coefficients between alternative allele frequencies (AAF) obtained from lofreq for WGS and rRNA-seq, with and without accounting for methylation levels, calculated on mouse kidney data from six different strains: 129S1/SvImJ (‘129S’), C3H/HeJ (‘C3H’), C57BL/6J (‘BL6J’), C57BL/6N (‘BL6N’), CAST/EiJ (‘CAST’) and A/J (‘AJ’).

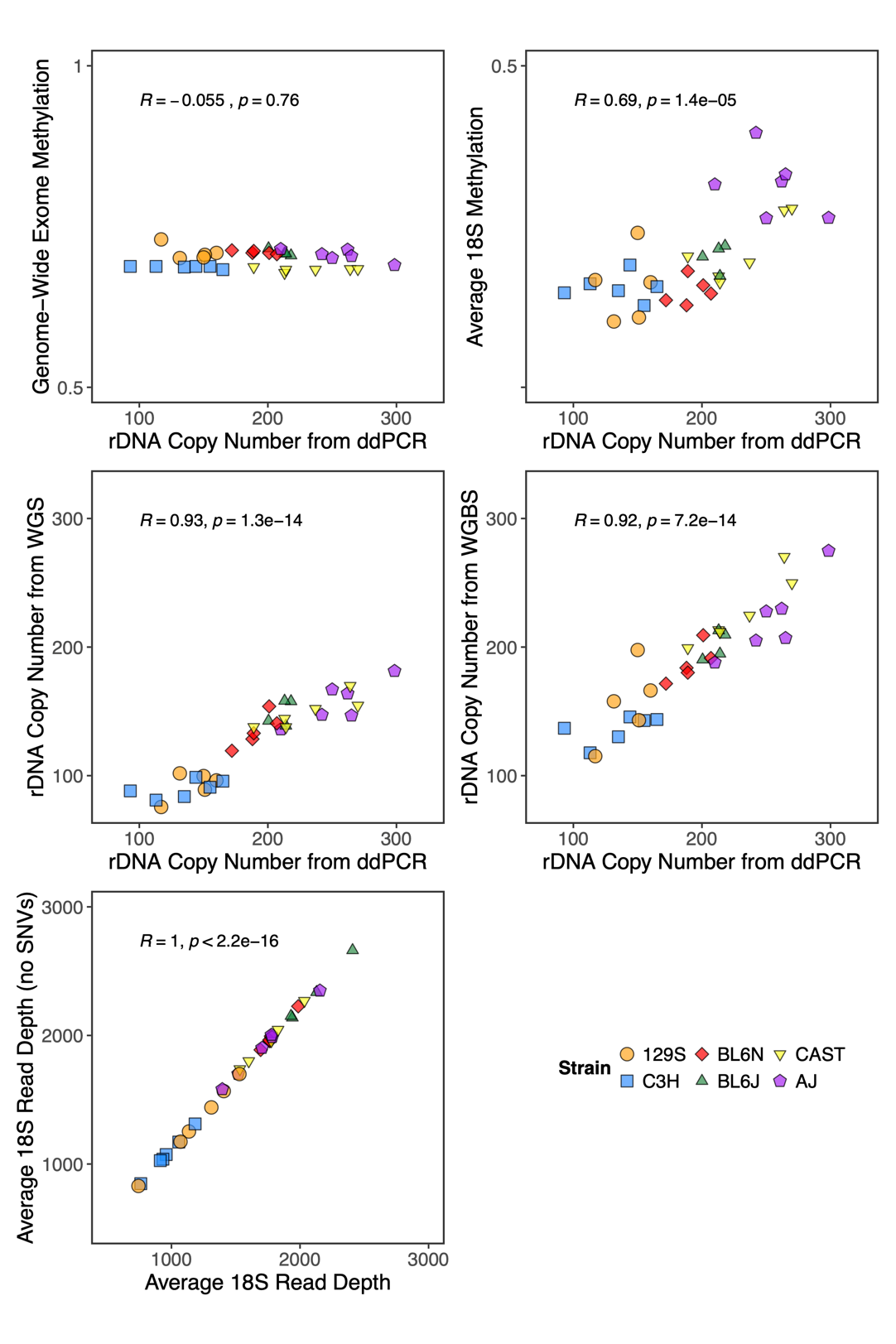

### **Figure S15. Association between ddPCR rDNA copy number with methylation levels and sequencing-based copy number estimates, on kidney mouse data.** Overall CpG methylation estimates from genome-wide exome alignments show no correlation with rDNA copy number (**Top left**), unlike CpG methylation at the 18S rDNA subunit (**Top right**). rDNA copy number estimates from WGS (**Middle left**) and WGBS (**Middle right**) correlate with ddPCR estimates. Average WGBS 18S read depth calculated across the entire subunit or only between positions 4418 and 5417 (longest stretch without variants, plus/minus 150 bp at each side) perfectly correlate (**Bottom**). All data points are calculated on mouse kidney data from six different strains: 129S1/SvImJ (‘129S’), C3H/HeJ (‘C3H’), C57BL/6J (‘BL6J’), C57BL/6N (‘BL6N’), CAST/EiJ (‘CAST’) and A/J (‘AJ’).

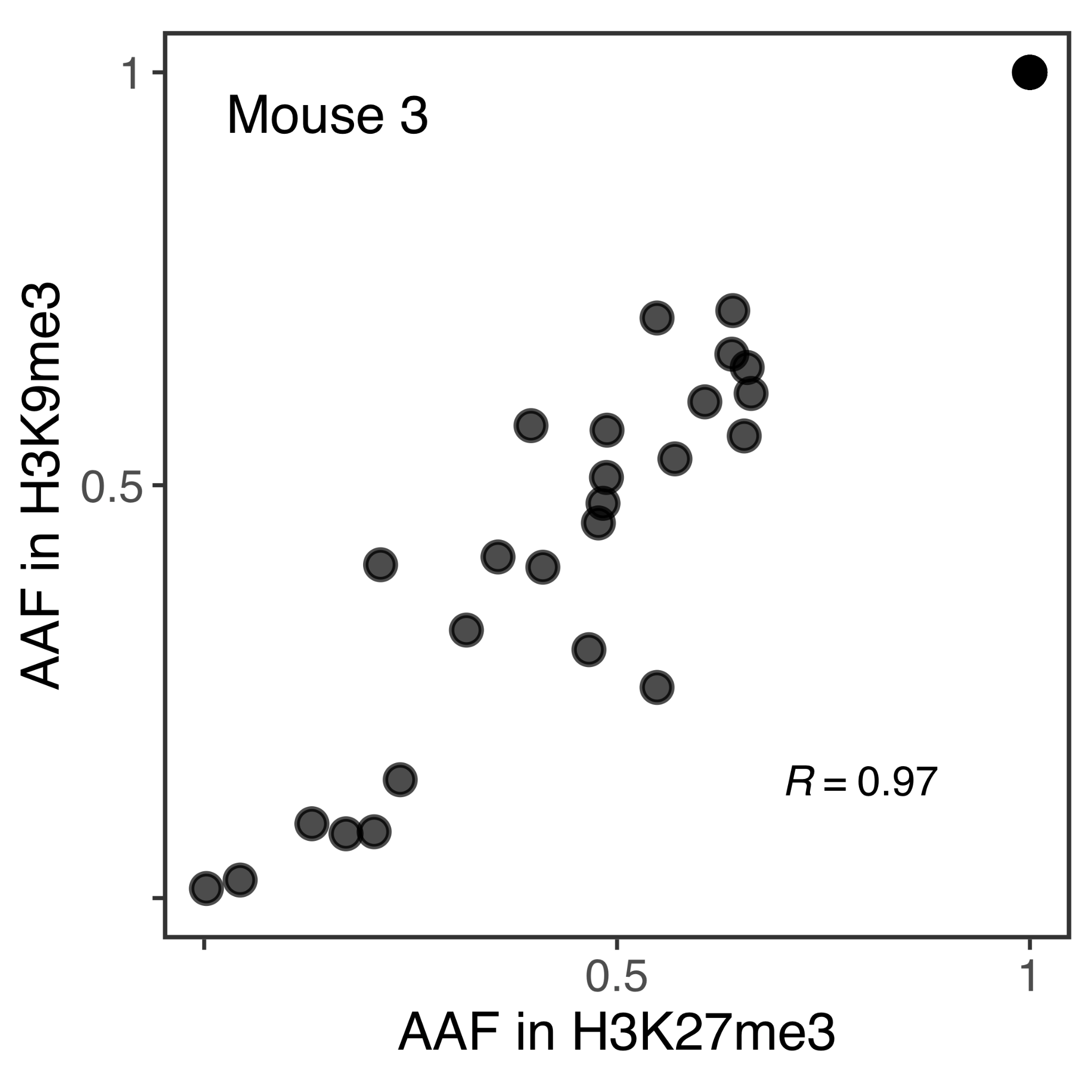

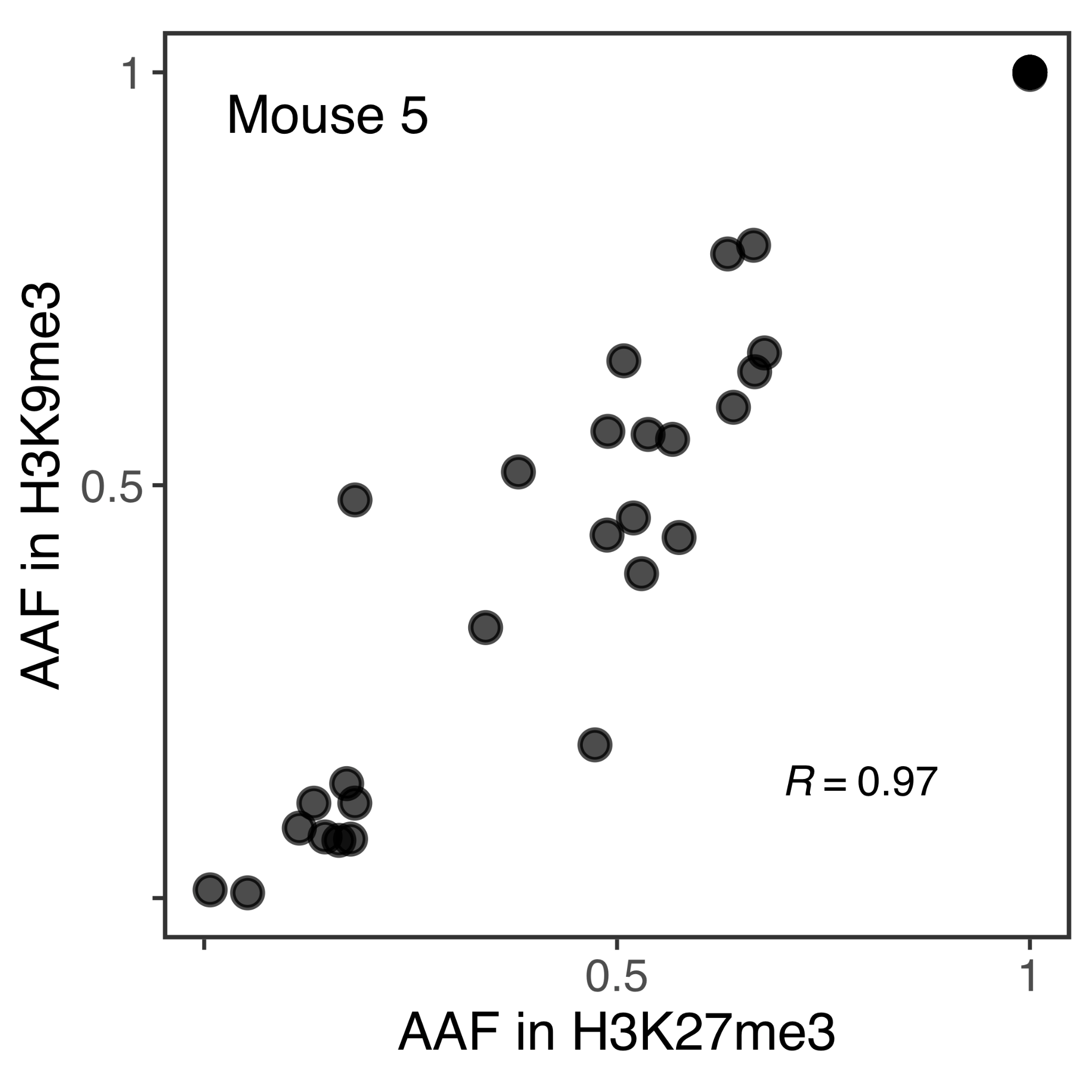

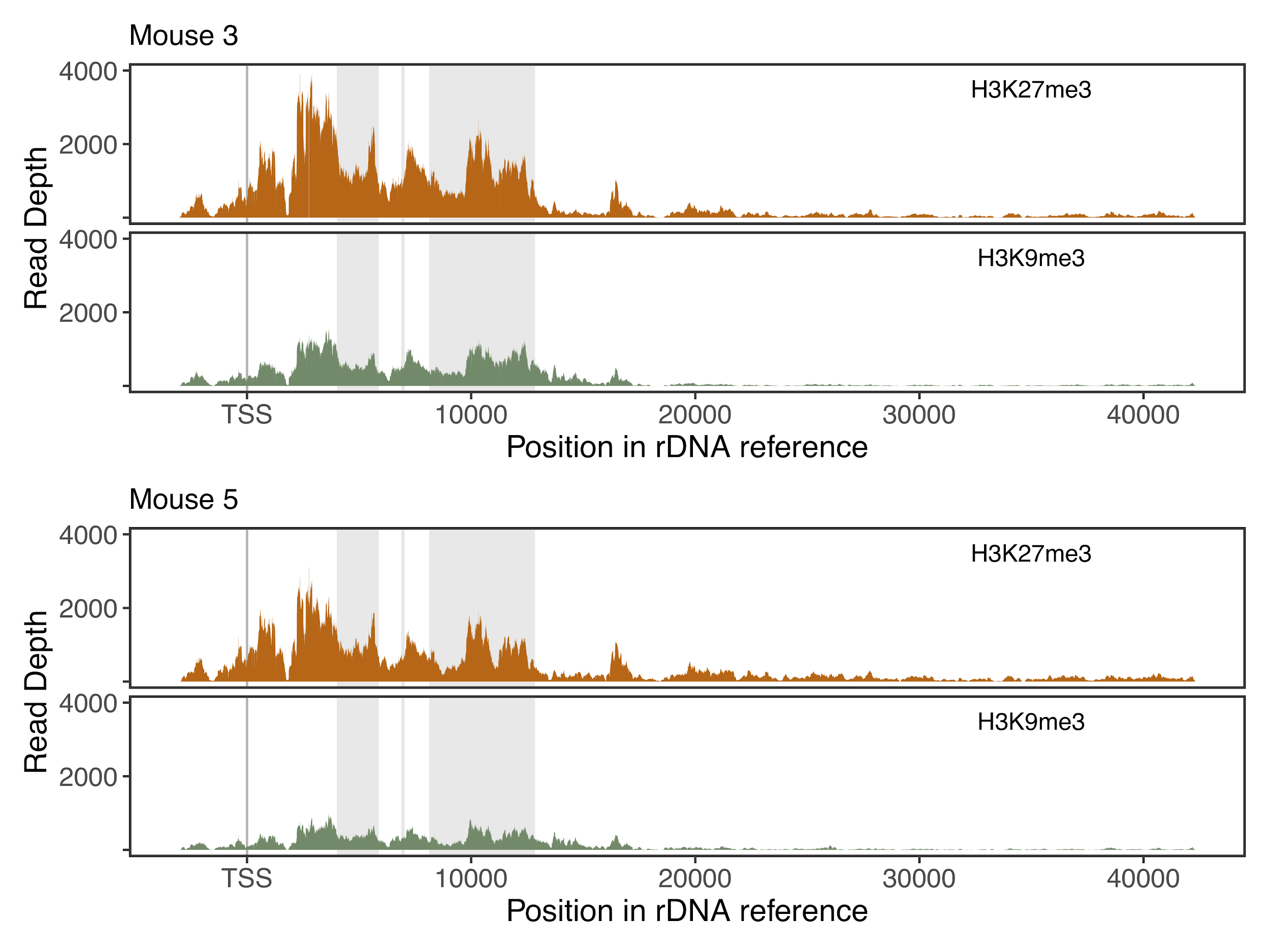

### **Figure S16. 129S1/SvImJ kidney CUT&Tag data.** Comparison of alternative allele frequencies at variant positions between H3K27me3 and H3K9me3 reads (**Top**), and read coverage distribution across the rDNA unit (**Bottom**).

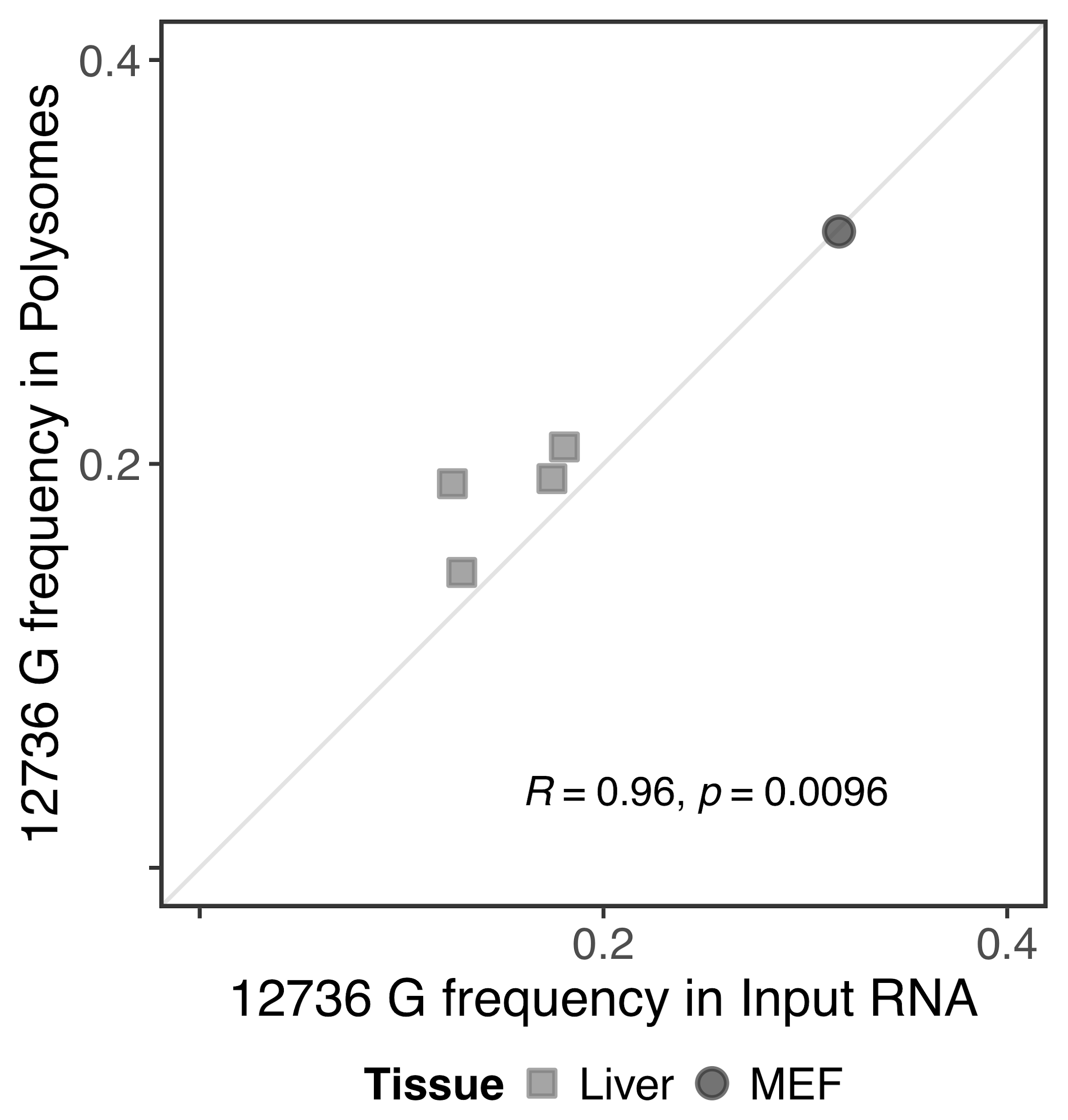

### **Figure S17. Association between and Polysomal RNA-seq allele frequencies.** Observed frequencies of the G allele at rDNA position 12736 in polysomes closely match those from total RNA-seq in C57BL/6J mouse liver and MEF data.

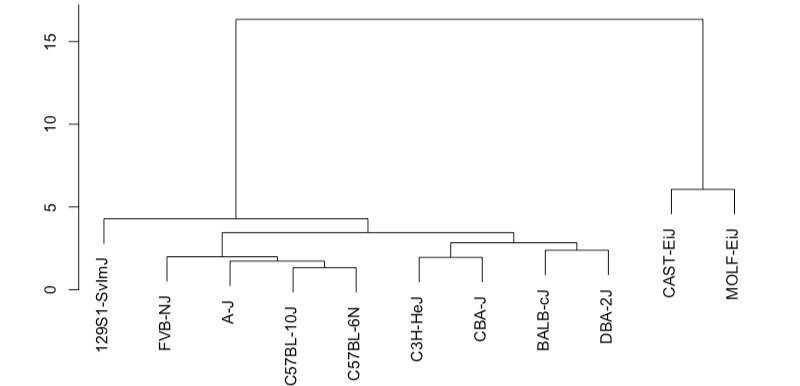

### **Figure S18. Clustering analysis of mouse strains according to rDNA variation.** Dendogram of the distance between mouse strains initially considered for the present study, calculated according to allele frequencies observed at rDNA variants.

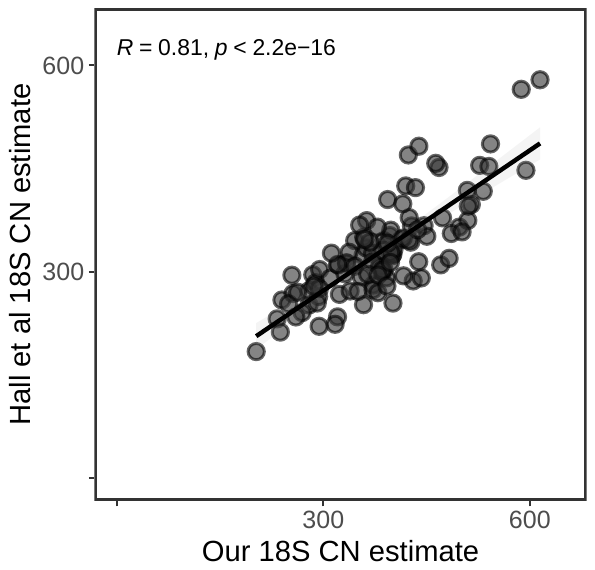

### **Figure S19. Correlation between publicly-available and our own rDNA copy number estimates for human Mandinka individuals.** Pearson correlation computed for the 108 human Mandinka individuals from the 1000 genomes project^32^ for which Hall *et al*^31^ provide rDNA copy number estimates from 18S.

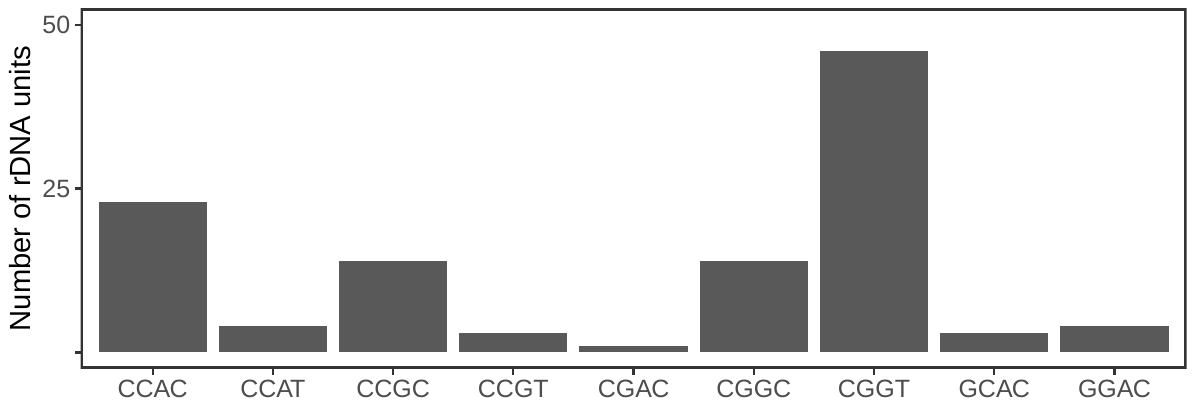

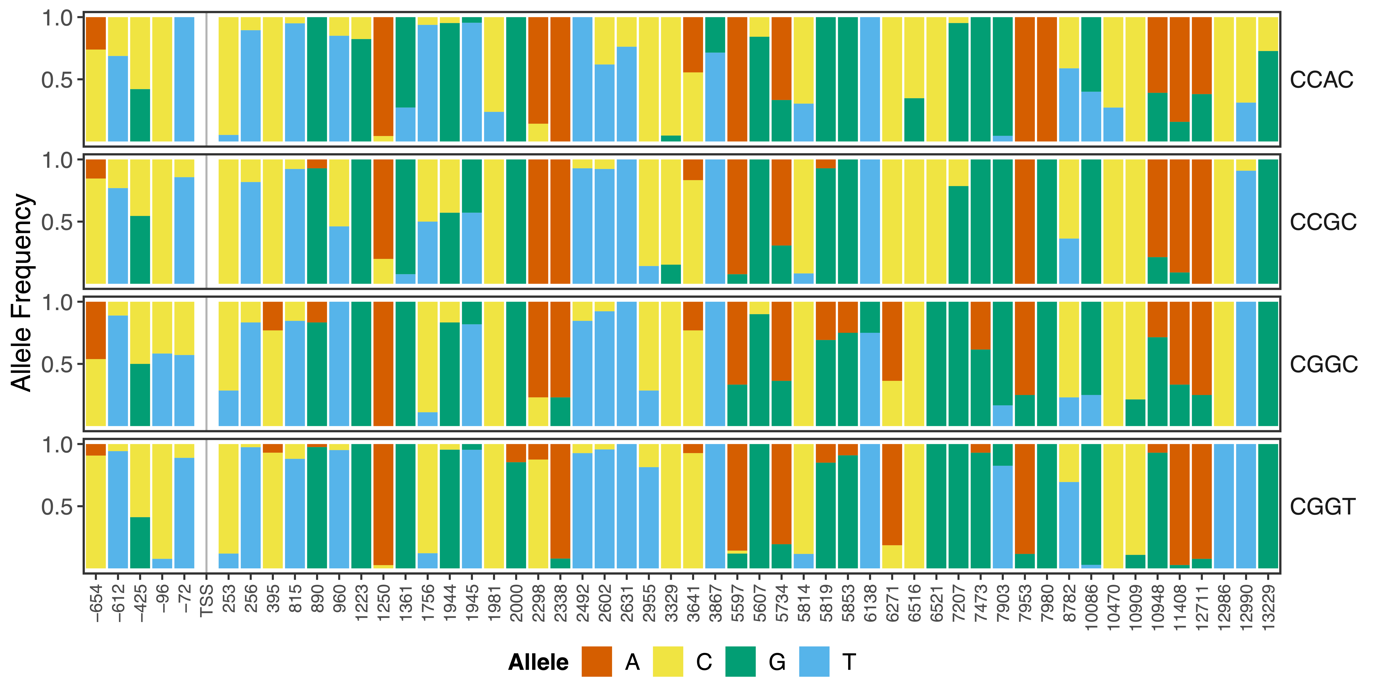

### **Figure S20. Example of putative haplotypic structure in a human Mandinka sample.** Number of rDNA units in the HG02723 ultra-long read Nanopore dataset for each observed combination of alleles at positions -348, 6521, 7980, and 12986 (**Top**), and allele frequencies at variant positions across the rDNA coding unit for the combinations with more than 5 associated units (**Bottom**).
